# Scalable Inference and Identifiability of Kinetic Parameters for Transcriptional Bursting from Single Cell Data

**DOI:** 10.1101/2025.05.13.653671

**Authors:** Junhao Gu, Nandor Laszik, Christopher E. Miles, Jun Allard, Timothy L. Downing, Elizabeth L. Read

## Abstract

**Motivation:** Stochastic gene expression and cell-to-cell heterogeneity have attracted increased interest in recent years, enabled by advances in single-cell measurement technologies. These studies are also increasingly complemented by quantitative biophysical modeling, often using the framework of stochastic biochemical kinetic models. However, inferring parameters for such models (that is, the kinetic rates of biochemical reactions) remains a technical and computational challenge, particularly doing so in a manner that can leverage high-throughput single-cell sequencing data.

**Results:** In this work, we develop a Chemical Master Equation reference library-based computational pipeline to infer kinetic parameters describing noisy mRNA distributions from single cell RNA sequencing data, using the commonly applied stochastic telegraph model. Our pipeline also serves as a tool for comprehensive analysis of parameter identifiability, in both *a priori* (studying model properties in the absence of data) and *a posteriori* (in the context of a particular dataset) use cases. The pipeline can perform both of these tasks, i.e. inference and identifiability analysis, in an efficient and scalable manner, and also serves to disentangle contributions to uncertainty in inferred parameters from experimental noise versus structural properties of the model. We found that for the telegraph model, the majority of the parameter space is not practically identifiable from single cell RNA sequencing data, and low experimental capture rates worsen the identifiability. Our methodological framework could be extended to other data types in the fitting of small biochemical network models.

**Availability:** All code relevant to this work is available at https://github.com/Read-Lab-UCI/TelegraphLikelihoodInfer.

## Introduction

When experiments on gene expression in cells reached single molecule resolution, it was discovered that the fundamental cellular process of transcription is surprisingly noisy (reviewed in [1]). Temporal measurements of individual transcribed mRNA molecules revealed that transcription occurs not smoothly and continuously, but in bursts [2]. So-called “intrinsic” biochemical noise, arising from the inherent stochasticity of biochemical reactions occurring in low copy number regimes, can partially account for transcriptional noise [3]. More broadly, important roles for biochemical noise have been identified in numerous cellular processes, including cell fate diversification in development [4–7], cellular reprogramming [8], cancer phenotype switching [9, 10], and bacterial antibiotic resistance [11].

Discrete, stochastic chemical reaction models have formed the basis of biophysical theories of gene expression noise [12–14]. The two-state gene expression model (also known as the telegraph model) has been a common baseline framework for understanding gene expression noise. In this model, the promoter of interest stochastically switches between two distinct ‘on/off’ activity states. The switching process could represent transcription factor binding/unbinding [15], mechanical forces [16], large-scale chromatin rearrangement occurring on slower timescales [17], etc.. Despite its simplicity, this model has gained traction due to its ability to recapitulate properties of gene expression noise from microbes to mammals, including heavy-tailed transcript distributions with negative binomial shape [18], and the scaling properties of noise with mean gene expression [19, 20].

Inference of kinetic parameters of the telegraph model from experimental data has been undertaken in a number of studies. Many of these involved fitting model-output distributions to experimentally measured mRNA histograms from single molecule fluorescence in situ hybridization (smFISH) experiments, either from single-timepoint snapshots [8, 21–25], or temporal measurements [19]. Recently, there has also been progress in inferring bursting kinetics from single cell RNA sequencing data (scRNAseq)[26–32]. The underlying concept is that of utilizing fluctuations, as measured across a population at steady-state, to infer dynamics. scRNAseq generates single-cell distributions for the entire transcriptome, in principle enabling comprehensive, genome-wide analysis of bursting kinetics. However, scRNAseq also suffers from significant technical noise [33], which makes parameter inference challenging.

Parameter identifiability is a subject that has received increased attention in recent years, including in the area of biochemical reaction modeling [34]. In general, parameters are considered structurally identifiable when a parameter set maps to a unique model output [35, 36]. In the case of unlimited, perfect data, the inference and identifiability problem are dependent on the structure of the physical model itself. “Structural identifiability” is distinguished from “practical identifiability”, which can be caused by insufficient data, loss of information, or noise sources affecting measurements, all of which could lead to unacceptably large uncertainty of parameter estimates. In model-guided analysis of scRNAseq, practical identifiability (the focus of this study) is of interest because of technical errors and noise in the measurement technologies, and because the sample size is often limited.

Researchers have leveraged various modeling and inference tools to infer kinetic parameters of the telegraph model from scRNAseq, and many of these have included some type of uncertainty analysis [26, 28–30, 32]. However, uncertainty analysis is generally performed in an *a posteriori* manner (i.e., in the context of a particular data set, after the fact)[34]. In this study, we sought to address the question of whether the parameters of the telegraph model are fundamentally, practically identifiable from scRNAseq data in general, rather than proceeding from the assumption that they *are* identifiable. To this end, we developed a computationally efficient batched Chemical Master Equation-based pipeline that performs both *a priori* and *a posteriori* inference and analysis of parameter identifiability. We found that, in most of the biophysical parameter space for the telegraph model, the parameters are not practically identifiable from snapshot mRNA distributions alone, even with 100% capture rate and large sample size. We assess statistical noise measures–i.e., distribution shape features–that can partly, but not fully, predict the identifiability of kinetic parameters for a given gene. In general, based on the telegraph model, only genes undergoing slow promoter switching have identifiable kinetics. Our pipeline and conceptual framework could be applied to other types of small stochastic biochemical network models and measurement contexts, or adapted to more complex models in the future.

## Methods

### Telegraph model and Chemical Master Equation solution

The reactions of the telegraph model are as follows:

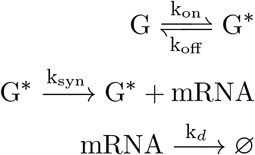

where *G*^*\**^, *G* represent the states of the promoter of the gene of interest (active, inactive, respectively). The gene state switches between active and inactive with reaction rates *k*_*on*_ and *k*_*off*_, respectively, and mRNA is produced at rate *k*_*syn*_ only in the active state (Fig. 1A). *k*_*d*_ is the mRNA degradation rate. We take the inverse degradation rate to be unit time of the system, thus setting *k*_*d*_ = 1. Kinetics of the telegraph model are sometimes characterized in terms of transcript burst size *k*_*syn*_*/k* and burst frequency 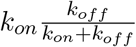.

**Figure 1:**
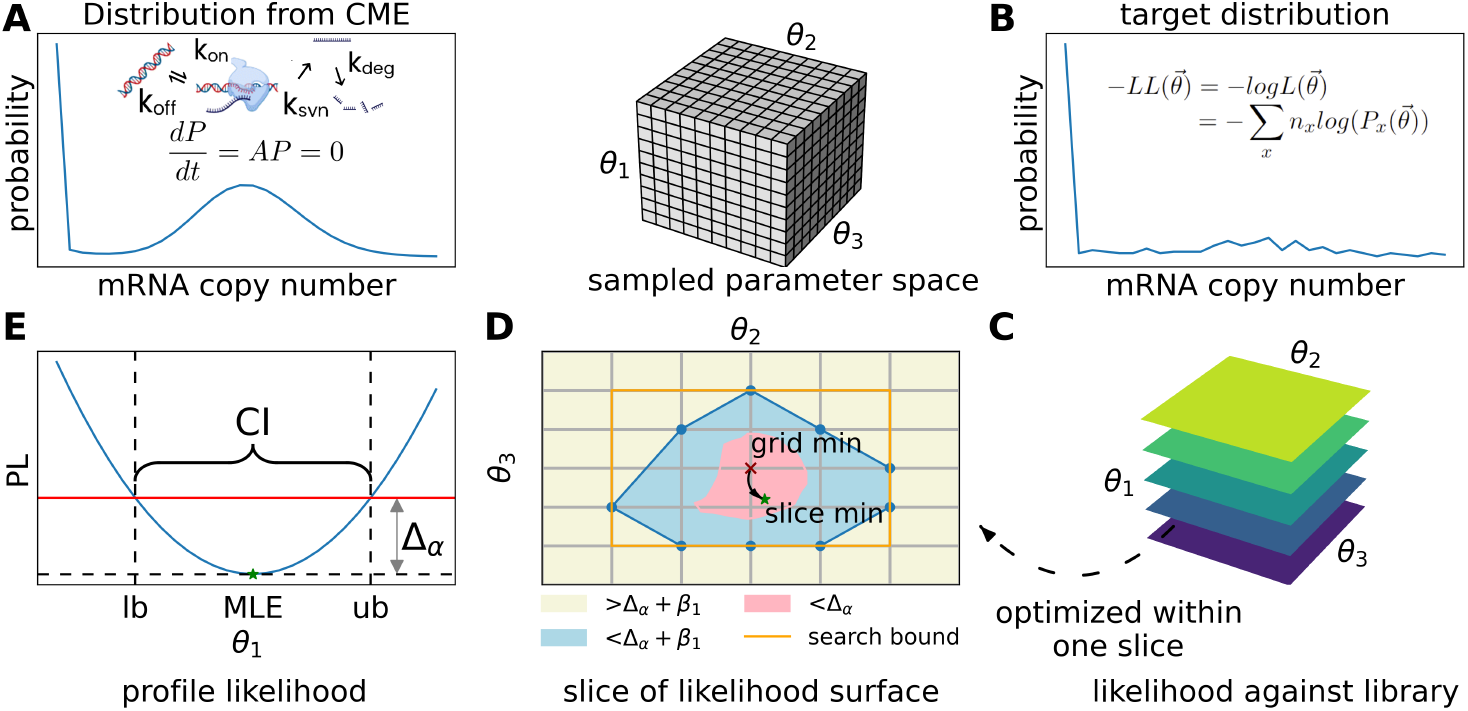
Computational pipeline workflow. a) mRNA distribution computed from the telegraph model, where the promoter switches between inactive and active states. Parameter sets are sampled as a 3-dimensional grid library for the parameters *k*_*syn*_, *k*_*off*_, and *k*_*on*_ (see Methods). b) Representative experimentally measured target distribution, from which the negative log likelihood (-LL) of sampled parameter sets 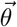 can be obtained by comparison to computed distributions. Alternatively, the target distribution can be obtained from synthetic data (i.e., model-generated distributions) for *a priori* identifiability analysis. c) The coarse-grained, 3D surface, i.e., the -LL value of every simulated mRNA distribution from the library against the target distribution. d) A schematic slice from the 3D -LL surface, demonstrating the optimization procedure: optimization is only performed within the search bounds obtained from the initially sampled coarse-grained -LL surface e) After optimization, the profile likelihood (PL) function for each parameter is obtained and confidence intervals are computed (see Methods and SI 1).

Although the telegraph model admits an analytical solution [12], we adopt the Chemical Master Equation (CME) framework because it can also be applied to other models. The system can be expressed in vector-matrix form by enumerating a finite number of states (i.e., neglecting low probability states) The CME is expressed as

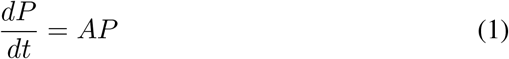

where 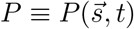 is the probability to find the system in system state 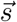 at time *t*, and *A* is the reaction rate matrix, whose elements *A*_*ij*_ give the rate of the reaction bringing the system from state *j* to state *i*, given by the model’s kinetic parameters and standard chemical rate laws. The steady state solution 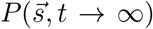 is thus obtained from *AP* = 0.

For convenience, we hereon refer to the steady-state model solution on the full state space as 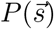. This solution can be further projected onto the mRNA copy number axis *x*, giving *P*(*x*), by summing over the on/off promoter states. This is done because promoter state is not directly distinguishable from scRNAseq data.

Scaling by *k*_*d*_, there are three inference parameters, as vector 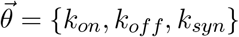. We hereon refer to the model-computed steady state distribution over mRNA molecules, given parameter vector 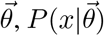 as 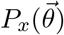, or 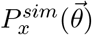, to further distinguish from an experimentally measured target distribution.

### Parameter estimates and confidence intervals

We use a standard approach for *a posteriori* parameter inference, using Maximum Likelihood Estimation (MLE) and Profile Likelihood (PL) functions to derive Confidence Intervals (CI) (Fig. 1E). For comparison of the model output to mRNA count distributions from scRNAseq, the log-likelihood function over parameter sets 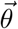 is:

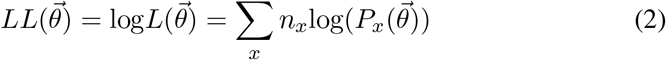

where *x* is the mRNA copy number and *n*_*x*_ is the number of cells with *x* observed mRNA copies (Fig. 1B). (Note: we use minimization of 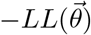 to obtain MLE).

### Profile Likelihood for *a priori* identifiability analysis

We first study the identifiability of the telegraph model in the idealized scenario: when the model itself is used to generate synthetic data. This is an *a priori* approach, because it depends only on the properties of the model itself and not on any particular dataset. For a hypothetical experiment with *N* cells and parameters 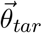, Eqn. 2 becomes

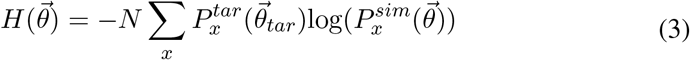

where 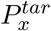 is the model-calculated probability of observing *x* mRNA in the target distribution, and 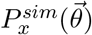 is the simulated probability of observing *x* mRNAs, given any parameter set 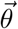.

For consistency with the *a posteriori* analysis and standard terminology, we also refer to the surface 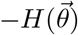 as the “log-likelihood surface,” while noting it is technically a scaled negative cross-entropy. To study identifiability *a priori*, the question is what shape 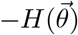 has. In the ideal case, it is narrowly peaked, yielding relatively narrow CIs. However, if varying values of 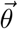 produce similar *P*, then the surface may have a broad peak with no clear global minimum–thus practically unidentifiable. The utility of Eqn. 3 is that one can study the effect of the cell number, *N*, in the hypothetical experiment without any error introduced by sampling.

We adapted the PL method to *a priori* analysis, whereas it is typically used in an *a posteriori* manner [34], by removing the need to use sampled data via Eqn. 3 (see SI 1.3). We refer to the PL so obtained as the “ground-truth PL”, equivalent to the average of sample-replicate-derived PL functions from infinite hypothetical experiments with *N* cells.

### Computational pipeline

#### Strategy to combine coarse-grained library with fine-grained optimization

Obtaining accurate MLE and CI involves non-linear optimization, which can suffer from local minima, early termination, and lack of efficiency, if poor initialization and boundary conditions are given. For a scalable approach (that deals efficiently with large numbers of experimental distributions, e.g. from transcriptome-wide data), we use a coarse-grained, simulated library of CME-derived-distributions as a reference. This provides coarse-grained estimates of MLE, PL, and CI, and also provides reasonable initial guesses and boundary conditions for further optimization (Fig. 1D and SI 1.4).

#### Generation of the reference library from the CME

We solved the CME with a grid sampling of the parameter space. For the three parameters, we took a resolution of 60 grid points for each parameter: *k*_*syn*_ : [10^*−*0.3^, 10^2.3^], *k*_*off*_ : [10^*−*3^, 10^3^], *k*_*on*_ : [10^*−*3^, 10^3^], (all in units of 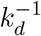) based on typical measured ranges [38, 39]. Using Eqn. 3, each target parameter set (i.e., the ground truth to be inferred) is thus mapped to a value of the *LL* surface at 60^3^ grid points (see Fig. 1C). The same grid points are also used as targets, thus the library entails (60^3^)^2^ precomputed *LL* values.

#### Profile Likelihood-based identifiability metric

We devised a PL-based metric for identifiability, termed the Alternative Precision Measure (APM):

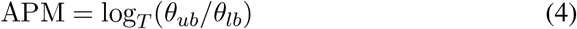

where *θ*_*ub*_ and *θ*_*lb*_ are the upper bound and lower bound of the CI, respectively, and *T* is user-defined. In general, the smaller the APM, the smaller the CI and the more identifiable the parameter. We apply a simple cutoff: when APM *<* 1, we consider the model (at that point in parameter space) to be practically identifiable. An APM below 1 indicates that the CI ratio falls within the chosen factor *T*. These threshold values (*T*) define a practical criterion for classifying identifiability, based on typical parameter scales and the sensitivity of the model. While the precise cutoff is subjective, overall trends in identifiability remain robust across parameter space. We used *T* values {*k*_*syn*_, *k*_*on*_, *k*_*off*_} = {3, 100, 100}.

To obtain a single APM value at a given point in parameter space, we use the maximum of the three individual parameter APM values, reasoning that the largest value reflects the “worst” parameter, i.e., the one that is least identifiable from the data. (Note this is similar to a related, recent method that used the union of profile-wise prediction intervals [40].)

We also studied the more common Precision metric for comparison, Fig. S1 and SI 2.1.

## Results

### Representative cases; Profile Likelihood matches bootstrapping MLEs

We demonstrate the computational approach with two representative parameter scenarios (Fig. 2), one of which is practically identifiable, and one which is not. In each case, the ground truth distribution is shown together with the *LL* surface over the parameter space and the recovered PL in the three separate parameter dimensions. (For ease of visualization, we project the 3D LL surface onto the 2D space of burst frequency and burst size). Additionally, we present results of sampling replicate synthetic distributions, which enables estimation of CIs from bootstrapping.

**Figure 2:**
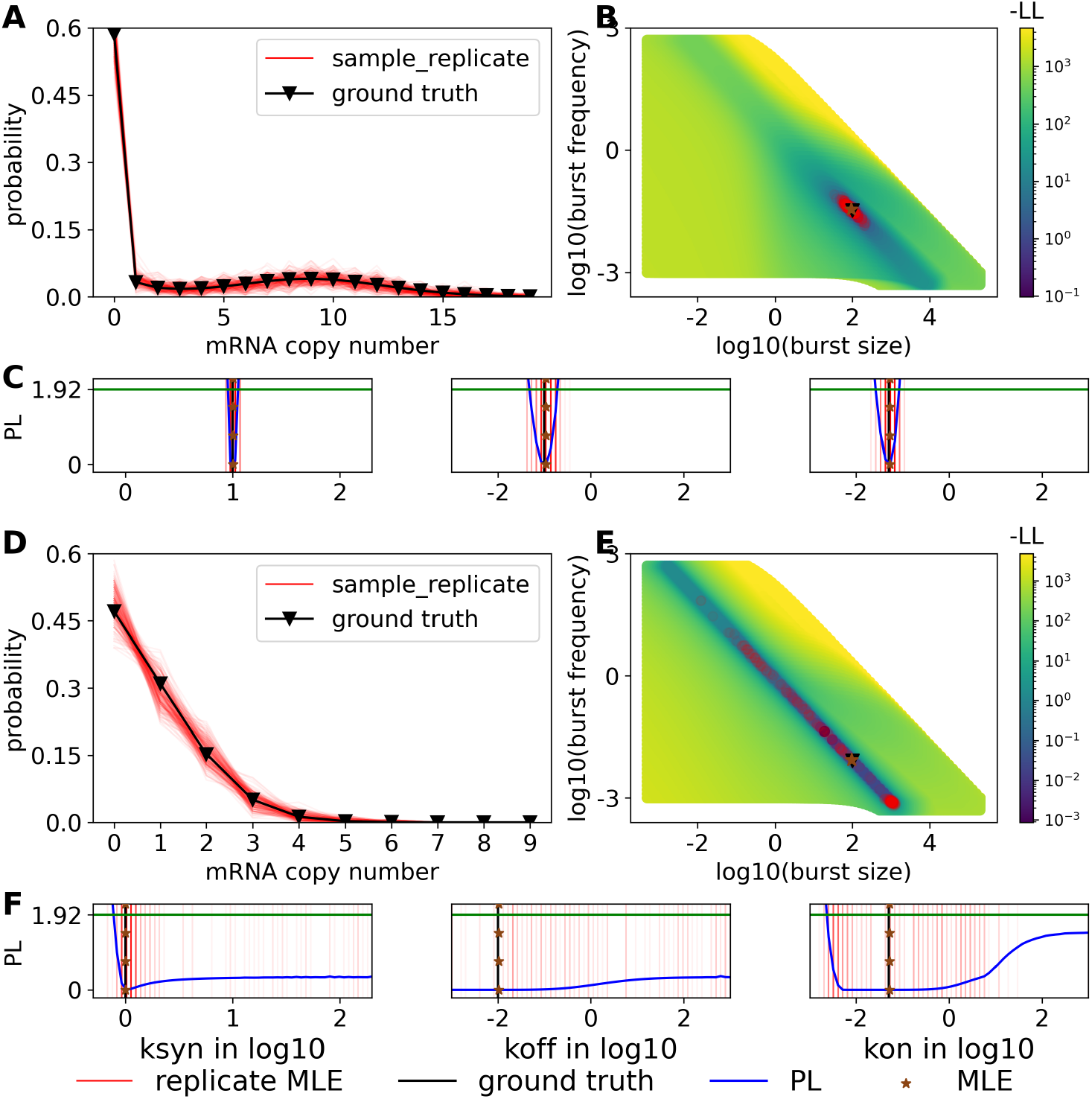
Profile likelihoods (PL) from two representative parameter sets with 200 cells: Panels (A,B,C): Representative parameter set that is identifiable, [ksyn:10, koff:0.1, kon: 0.05], (D,E,F): Representative parameter set that is practically unidentifiable, [ksyn:10, koff:0.01, kon:0.05]. A, D) Original computed distributions (black), and sample replicates (red); B, E) 3D -LL surface projected onto 2D burst frequency and burst size (performed as a scatter plot of sorted -LL values in 2D; when there is overlap, smaller values are in front); C, F) PL of the three parameters: *k*_*syn*_, *k*_*off*_, *k*_*on*_. For the red dots and stripes, the intensity indicates the frequency of the replicate MLE. The overall PL distribution covers the parameter range where the MLEs take place. The green horizontal lines indicate the 1.92 *χ*^2^ value.

For the identifiable case (Fig. 2A-C), PL functions are narrow and MLE replicates are narrowly distributed for each parameter. In contrast, in the practically un-identifiable case (Fig. 2 D-F), the PL is so broad that either a global minimum is not found, or if found, a majority of the parameter space lies within the confidence region. These scenarios demonstrate how the parameters of the telegraph model may or may not be identifiable, depending on the kinetic regime and sample size.

We observe consistency between the distribution of the sampled MLEs (red) and the ground truth PL (blue) (Fig. 2C,F), indicating also consistency between the CIs inferred from both methods. The advantage of our *a priori* approach is that it accounts for the loss of identifiability because of finite cells, yet does not contain any error due to sampling (since it represents the limit of finite cells but infinite experiments). Our approach also has the advantage of increasing computational efficiency, since the cell number *N* is simply a scalar multiple of LL and PL. Thus one does not need to recompute the library or perform sampling to assess the impact of experimental cell number on identifiability.

### Most of the parameter space is un-identifiable

To assess identifiability, we comprehensively computed CIs, and hence APMs, across the parameter space. Fig. 3 (B,C) shows the APM (each parameter and combined) for two different experimental RNA capture rates (100%, 30%), over the entire 3D parameter space, for a sample size of 10^4^ cells (a realistic *N* value). However, capture rates typically range from 10-30%, depending on toolkits used[41– 44], so these results represent an optimistic scenario. Even so, in Fig. 3, for most of the explored parameter space, the model is not identifiable.

**Figure 3:**
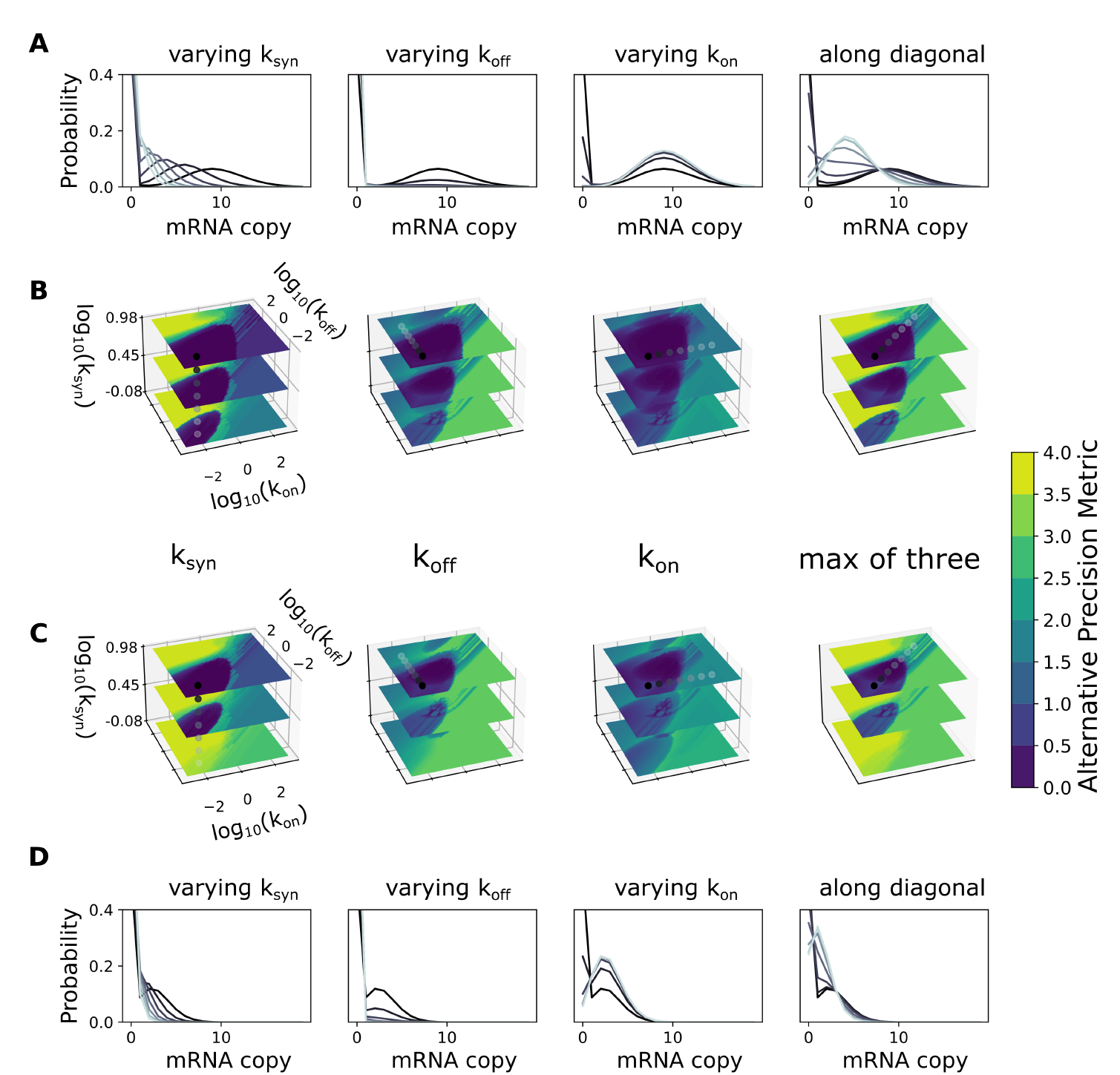
Global *a priori* identifiability landscape over the entire studied parameter space at different capture rates for 10K cells. (A,B) Results for 100% capture rate. (A) mRNA distributions for representative parameter sets. (B) Identifiability (measured by APM) at each ground-truth point in the 3D parameter space of each parameter separately (left three columns) and the overall identifiability (last column, maximum APM from all parameters). Distributions in (A) correspond to dots (grayscale color) in the corresponding 3D surfaces in (B). (C,D) Same as top rows, but with 0.3 experimental capture rate.

We observed that the region of parameter space showing practical identifiability is shaped like an inverted cone near the slow *k*_*on*_ and *k*_*off*_ regime, indicating that transcriptional bursting kinetics are most identifiable in the slow-gene-switching scenario. The corresponding distributions typically show bimodality at 100% capture rate, as seen in the distributions in darker color in Fig. 3A. Distributions (in lighter color) corresponding to either fast *k*_*on*_ or *k*_*off*_ are more Poisson-like and less identifiable. Note that with smaller *k*_*syn*_, the second mode is at low mRNA count, and only very slow *k*_*on*_ and *k*_*off*_ can practically lead to separation of the two modes and hence identifiability. These results of *a priori* analysis demonstrate that one should be careful with the assumption of identifiability for the telegraph model.

We also studied identifiability as a function of bursting kinetics. Burst-like transcription refers to fluctuations in mRNA synthesis that can be characterized by rapid synthesis (i.e., bursts) followed by periods of relative inactivity, and it has been observed from prokaryotes to mammals [2, 21, 45]. Transcription burst-size (i.e., average number of mRNAs produced in a single burst) and burst-frequency are often used to characterize kinetics [25, 28, 46], and these measures may be, arguably, of more interest from a biological standpoint than the kinetic rate parameters themselves. These measures can be directly obtained from parameters in the telegraph model (see Methods).

However, we noticed that this effective projection of the 3D parameter space *k*_*on*_, *k*_*off*_, *k*_*syn*_ onto 2D (burst-size and burst-frequency) can be misleading. Systems with the same burst-size and burst-frequency can differ greatly in identifiability (Fig. S2). Moreover, much of the (2D) parameter space is again unidentifiable according to APM. Therefore, inference of burst-size and burst-frequency from scRNAseq via the telegraph model may not be reliable.

### Effect of cell number and RNA capture rate

We explore the effect of capture rate and sample size on identifiability across a range of possible bursting kinetics. We characterize percentage of total parameter sets (from the 60^3^-size library) that are identifiable for capture rates ranging from 30-100% (i.e., at or better than current scRNAseq technologies) and cell numbers ranging from 100 to 10^8^. Across this range of cell numbers, to achieve the same percentage of identifiable parameter sets at 30% as compared to 100% capture, would require an increase of 10 times the number of cells, on average. Conversely, for a certain *N*, where *N* ≤ 10^4^, the percentage of identifiable parameter sets is reduced by at least half. The lower *N*, the more drastic the reduction of size of the identifiable parameter space (Fig. 4A). The effect of these technical noise sources on identifiability for specific genes depends on their underlying bursting kinetics. In a scenario with moderately slow switching between promoter states (Fig. 4B-I), approximately 100 times more cells are needed to achieve the same level of precision at 30% capture rate, as compared to 100%.

**Figure 4:**
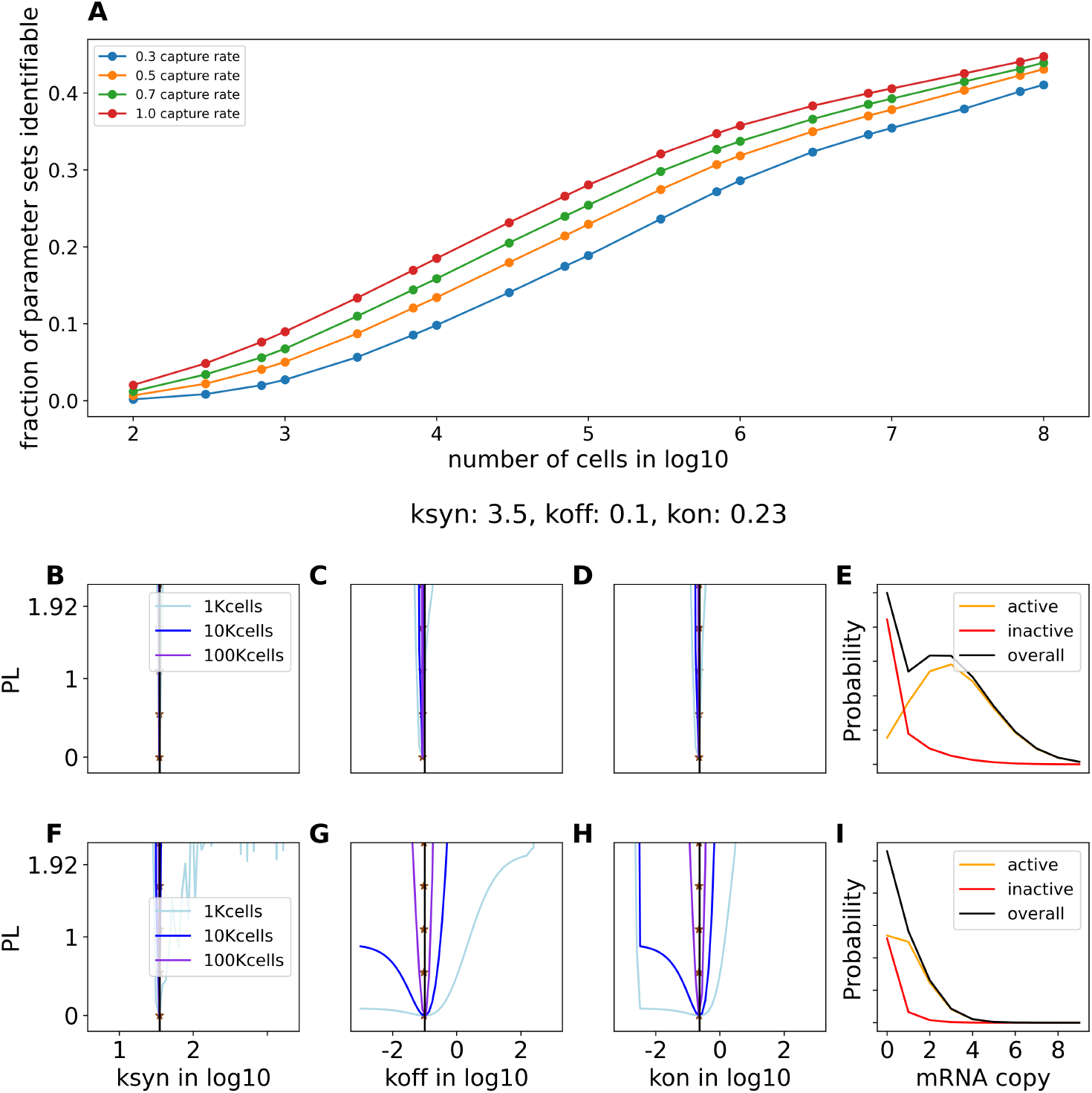
The effect of cell number and capture rate. (A) Fraction of identifiable parameter sets from the whole library vs. number of cells at different capture rates; (B-D) the Profile Likelihoods for a representative parameter set {*k*_*syn*_ : 3.5, *k*_*off*_ : 0.1, *k*_*on*_ : 0.23} at 100% capture rate, cell number 1K (light blue), 10K (blue), 100K (violet). (E) The mRNA distribution conditioned on active (G^*\**^) and inactive (G) promoter states. (F-I) Corresponding results for the same parameter set as to (B-E) for 30% capture rate.

Bimodality is a key feature of the telegraph model, which is linked to the effects of both *N* and capture rate on identifiability. The bimodality arises from separation between mRNA distribution peaks in the two promoter states, G, G^*\**^, which occurs when *k*_*on*_ and *k*_*off*_ take low or moderate values. This is seen by computing the mRNA distributions conditioned on each of the two promoter states (Fig. 4E,I). For the scenario in Fig. 4B-E, at 100% capture rate there is clear separation between the modes, resulting in narrow CIs over all parameters, even at the smallest sample size. For the same parameters, mode separation is lost due to signal degradation at lower capture rate (Fig. 4F-I), thus requiring at least 100K cells to achieve identifiability of all parameters. The problem is exacerbated for genes with low transcription rates (*k*_*syn*_), because one can less afford signal degradation when few molecules are present to begin with. Thus, low-*k*_*syn*_ genes require especially slow switching kinetics (*k*_*on*_, *k*_*off*_) for identifiability. Conversely, high transcribing genes have identifiable switching kinetics over a broader range.

Another scenario where low capture rate is especially detrimental is one where gene activity is intermittent, i.e., long periods of inactivity punctured by brief periods of activity. This occurs when both *k*_*on*_ and *k*_*off*_ are slow or moderately slow, and *k*_*on*_ *< k*_*off*_, resulting in bimodality with low probability in the active mode. Here, the model sometimes fails to distinguish between a bimodal distribution (with low probability in the high peak) and a corresponding unimodal distribution.

### Partial consistency between Profile Likelihood and other measures for *a priori* identifiability analysis

To validate our findings on the global identifiability landscape based on PL (i.e., in Fig. 3), we compare the results to those obtained from other approaches. One can calculate the sensitivity of the model output to parameter changes directly (i.e., without data), hence sensitivity analysis is an *a priori* approach. The elements of the sensitivity matrix 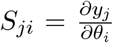 give the change of model output *y*_*j*_ with respect to parameter *θ*_*i*_. The magnitude of singular values of matrix *S* was suggested as a measure of parameter identifiability in dynamical systems[47]. Other studies utilized similar techniques based on the sensitivity matrix derived from the CME, such as the Fisher Information Matrix[48, 49].

Our sensitivity analysis, based on the minimum singular value of the sensitivity matrix (Fig. S3A), supports the telegraph model as structurally identifiable across the explored parameter space, as non-zero singular values were consistently found (albeit very small, 𝒪 (10^*−*15^) in some regions). This indicates that the widespread lack of identifiability observed (Fig. 3) is overwhelmingly a *practical* rather than *structural* issue, arising from low model sensitivity combined with finite sample sizes and experimental noise.

The minimum singular value, computed over the studied parameter space, qualitatively resembles the overall output of our PL-pipeline output (Fig. S3A). Specifically, the region of increased sensitivity occupies the slow promoter-switching kinetic region (low *k*_*off*_ and *k*_*off*_), and the size of this region increases with *k*_*syn*_. These results demonstrate that the PL is a viable way to access similar information to the sensitivity. However, our PL-based approach has the advantage that the effect of cell number and capture rate can be included in the analysis, rendering it a more pragmatic approach.

We further explored how various RNA-distribution summary statistics varied as a function of the parameter values, reasoning that it could be useful if simple summary statistics correlated with identifiability, thus providing an alternative means of *a priori* analysis. We studied various summary statistics based on moments of the distribution, including the the Fano factor *σ*^2^*/µ* (often used as a measure of dispersion in gene expression) (see Fig. S3 and S4). In general, measures related to second or third moments correlated, albeit only somewhat, with identifiability. For example, the Fano factor can be relatively insensitive to bimodality when one mode has very low probability (further details in SI 2.2). Thus, no single shape metric studied could replace the PL-based pipeline as a predictor of identifiability.

### *A posteriori* parameter inference from data: Few genes have identifiable bursting kinetics

We applied the PL-based inference pipeline to two scRNAseq datasets: SS3 cast of mouse fibroblast (CAST/EiJ × C57BL/6J)[28] and HUES64 human embryonic stem cells[50]. We found that only a small fraction of individual genes’ kinetic parameters are identifiable, according to our criterion (maximum APM from all parameters *<* 1). 970 gene distributions out of 10700 genes are considered to have identifiable kinetics for SS3 data, while 1153 genes out of 18806 are identifiable for HUES64WT (Fig. 5). As expected from the *a priori* analysis, genes for which parameters were inferred with high confidence tend to lie in regions of the parameter space with high expression rates and slow to moderate promoter switching kinetics. In summary, these results demonstrate that identifiability of transcriptional bursting kinetics from snapshot scRNAseq data is generally poor. Thus, these *a posteriori* results are in general agreement with the findings from *a priori* analysis.

**Figure 5:**
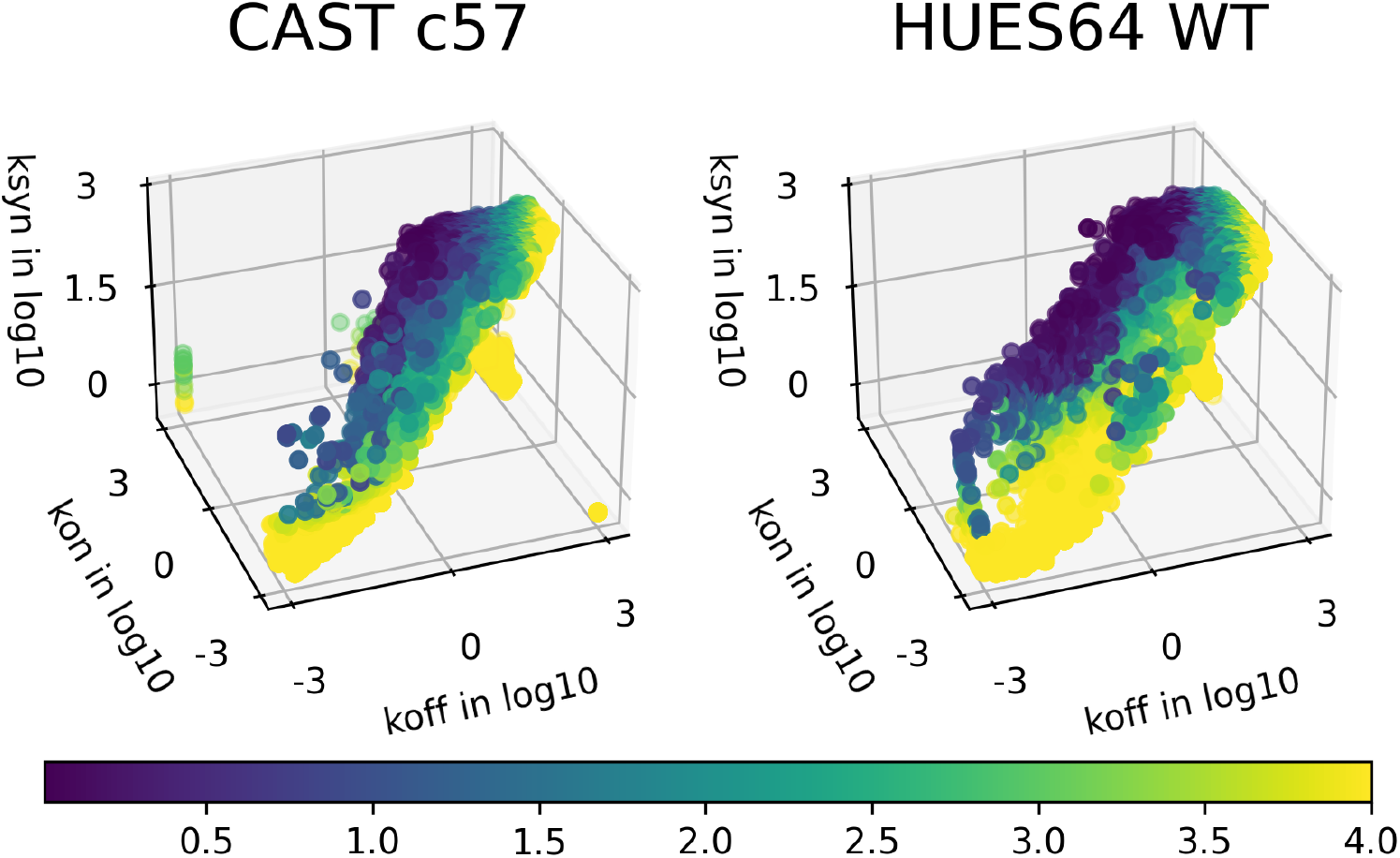
Inferred kinetic parameters of the telegraph model based on two datasets: SS3 cast of mouse fibroblast (CAST/EiJ × C57BL/6J) with cell numbers ranging from 6 to 224 (with mean 208), and HUES64 human embryonic stem cell with cell number of 1112. Each dot corresponds to a gene in the dataset. Color indicates the identifiability of the gene, as quantified by the APM metric derived from Profile Likelihood-based CIs (maximum over all three parameters). Only a small fraction of genes can be inferred with narrow CI, and thus reach the criterion of identifiability (APM*<* 1).

Since identifiability is related to distribution shape, we asked whether it would be possible to rapidly assess identifiability for a given experiment-derived mRNA distribution (together with information on cell number and capture rate) without applying the full PL-based inference pipeline. A neural network was trained on synthetic data to predict APM based on various distribution summary statistics (SI 2.3 and Fig. S5.) We then applied the neural network predictor to the real-world datasets, in order to assess its performance.

In general, there was good overlap between the set of identifiable genes (via PL) and those via the neural network. However, many genes were falsely predicted by the neural network to have identifiable kinetics. (Details and discussion of the false-prediction scenarios in SI 2.3). These results demonstrate that, for the real-world data as well as the synthetic data, combinations of distribution summary statistics (as leveraged by the neural network) can only partially predict the identifiability of the telegraph model parameters.

## Discussion

We developed a pipeline to infer kinetic parameters describing noisy gene expression from transcript distributions, such as those obtained from snapshot scRNAseq. Our approach differs from recent, related studies and methods [26–30, 32] in two key respects. First, we comprehensively analyze the practical identifiability of kinetic parameters of the classic telegraph model from single cell data, whereas many related studies report inferred kinetics without critically assessing the identifiability (or lack thereof) or parameters. Second, our pipeline is efficient and scalable: it integrates a batched CME solver with the PL method to quantify parameter uncertainty, utilizing a reference library of solutions to the CME model. This reduces reliance on optimization and redundant calculations, making the pipeline amenable to analysis of high throughput transcriptomic data. Another unique aspect of our approach is that we apply the PL method both to *a priori* investigation of identifiability and *a posteriori* inference of parameters; this ensures that the *a priori* study is more directly applicable to the practical output that researchers seek in conducting parameter inference. To this end, our pipeline also incorporates experimental parameters (cell number, capture rate) into the *a priori* analysis. We find that it is often more beneficial to increase capture rate, rather than cell number, to increase identifiability of parameters.

Our results show that the major part of the biologically feasible parameter space for the telegraph model is not practically identifiable from mRNA distributions, according to a criterion that we developed based on precision of inference. We confirmed the result using representative scRNAseq datasets. Sensitivity analysis showed that the model is structurally identifiable, even while practical identifiability is low (echoing previous results on the structural identifiability of a related telegraph model with protein translation [51]). In all, these findings underscore the potential pitfalls associated with attempting to infer complex stochastic dynamics from datasets with few degrees of freedom (as in, one dimensional distributions). Despite these potential pitfalls, our study also highlights how even 1D mRNA distributions can, in certain cases, have subtle but distinctive features that can be leveraged by the CME-based inference pipeline to elucidate underlying kinetics, especially in the slow-promoter-switching regime, which is associated with bistable transcript distributions. Identifiability of parameters in this regime has been established previously [26, 31]. We also find that “borderline” (not clearly bistable) cases can be identifiable (e.g., for moderate promoter switching, low transcription rate, and/or high dropout due to low capture rates in experiments). Such cases can introduce subtle but detectable shifts from Poisson, which can be utilized by the pipeline. Simple summary statistics, such as the commonly used Fano factor, often fail to recognize these borderline cases; as such, this argues for the telegraph model parameters as a more detailed list of shape features to describe noisy gene expression.

Our use of a CME solution library, for comparison of the model output to target distributions, is possible because the telegraph model is “small”, i.e. it is a biochemical network model with a limited feasible state-space and a small number of parameters, rendering both the size of the enumerated transition rate matrix, and the size of the feasible parameter space, to be tractable. A number of recent, related studies have combined Bayesian methods with stochastic simulation [52–60]. The advantage of our approach lies in the relative efficiency of CME solution for small models. Furthermore, the generated library can be used as reference to decide whether optimization is needed, in contrast to Bayesian approaches, where output distributions are calculated once the parameter prior is sampled, and discarded after the posterior is obtained. However, in use-cases with a small number of target distributions (i.e., few genes), and a more complex model (high-dimensional parameter space), using Bayesian-based methods may be more logical. Nevertheless our approach could be scaled to more complex biochemical models: as the dimension of model parameters increases, there comes a combinatorial increase of parameter sets. Keeping a manageable sized library would require a coarser parameter grid and thus more iterations to obtain CI.

The telegraph model has been widely used to describe gene expression noise, although many studies have also noted its shortcomings. It has been successful, for example, in describing statistical properties of transcript distributions [18–20] and elucidating the link between genomic features and gene expression noise [28]. Limitations of the model include its inability to describe biological mechanisms such as downstream processes [15, 61], feedback regulation [62], polymerase dynamics [63], multiple (*>* 2) promoter states [64], and more. The applicability (or not) of the telegraph model to describe real sources of noise in gene expression was not a focus of the present work. Instead, we focused on whether the parameters are identifiable, even in the idealized case, where the model perfectly describes the process to be inferred. Our results support a cautious application of the model to scRNAseq data: the parameters are often not identifiable, but may be for select cases. Our results suggest that more complex models will likely fail to be identifiable from the same type of data, since the issue of inferring multiple parameters from few-degree-of-freedom data would be even more pronounced. This also means that parameter identifiability should be carefully accounted for when comparing accuracy of different models to describe snapshot single cell data. Despite its shortcomings, the simplicity of the telegraph model may render it more appropriate for inferring noise properties from snapshot one-dimensional distributions than other models, whereas more complex (and biologically descriptive) models should be applied when additional data types are available.

Furthermore, establishing practical thresholds for parameter identifiability, such as our APM criterion, is crucial for downstream biological interpretation. Quantifying how transcriptional kinetics change in response to perturbations [65], or defining cell types based on underlying biophysical dynamics [66], requires parameter estimates with sufficient precision. Our analysis reveals the fundamental resolution limits imposed by the telegraph model when applied to snapshot mRNA distributions alone, suggesting that these data often lack the power to reliably distinguish subtle kinetic changes or classify states based solely on these parameters (i.e., APM is often *>* 1). Achieving the necessary parameter resolution for robust biological interpretation likely requires incorporating richer data types or spatial context that provide additional constraints. This could involve leveraging time-series measurements [2, 19, 62, 67], integrating multimodal single-cell data such as nascent and mature transcript counts [31, 66, 68], or accounting for the broader biological importance and specific modeling of mRNA spatial dynamics [69, 70]. These approaches move beyond the basic assumptions or data limitations explored here, highlighting the need for layered or spatially-resolved data to robustly connect bursting kinetics to cellular function.

## Acknowledgments

This work was supported by the following grants: NSF EF2022182, NSF DMS1763272, NSF CAREER DMS-2339241 and the Simons Foundation (594598).

## Supplementary Information to

### 1. Extended Methods

#### 1. Chemical Master Equation solution

We enumerate the Chemical Master Equation (CME) for the telegraph model as follows. Assuming that the number of mRNA molecules *x* is always 0 ≤ *x* ≤ *x*_*max*_, then the number of possible states of the system is *M* = 2 × (*x*_*max*_ + 1) (instantaneous state given by the promoter state (on or off) and number of mRNAs). We assume a value of 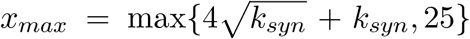. This *x*_*max*_ definition ensures low error (as the total probability of the truncated states is small) and is convenient for batch calculations. (More precisely, a gene at maximal activity (*k*_*on*_ ≫ *k*_*off*_) has Poisson-distributed expression with mean *k*_*syn*_, and the truncated probability under this definition is *<*10^*−*4^. For genes with lower activity, the truncated probability is lower.)

The CME is expressed as

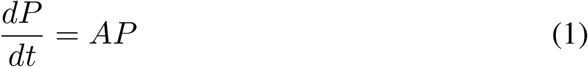

where 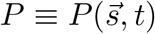 is the probability to find the system in system state 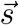 at time *t*, and *A* is the reaction rate matrix, whose elements *A*_*ij*_ give the rate of the reaction bringing the system from state *j* to state *i*, given by the model’s kinetic parameters and standard chemical rate laws. The steady state solution 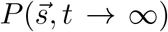 is thus obtained from *AP* = 0. Block representation is used to reduce the matrix size by half (for a detailed derivation, please refer to SI 3.1).

#### 2. Parameter estimates and confidence intervals

##### Maximum Likelihood Estimation

For comparison of the model output to mRNA count distributions from scRNAseq, the log-likelihood function over parameter sets 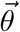 is:

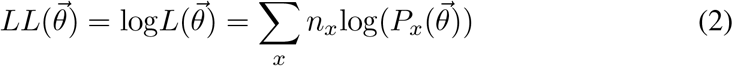

where *x* is the mRNA copy number and *n*_*x*_ is the number of cells with *x* observed mRNA copies (Fig. 1B). The maximum likelihood estimation (MLE) 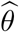 is then the infimum of Eqn. 2. We adhere to the use of minimization:

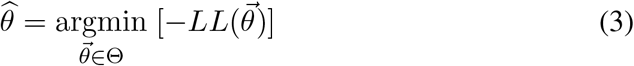

##### Profile Likelihood-based Confidence Intervals

Profile Likelihood (PL) is often used when accurate confidence interval estimates are difficult to obtain using standard methods, for instance, when *LL* is non-normal or when the model has high parameter dimensions[1]. PL is also useful for estimating confidence intervals (CI) for each parameter separately. Suppose *D* is sample data, and the fitted probability mass function is 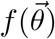. The parameter space can be partitioned as 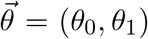, where *θ*_0_ is the parameter(s) of interest. The negative log-likelihood function (-LL) can be written as:

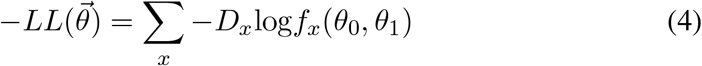

To get the profile likelihood of *θ*_0_, Eqn. 5 is evaluated :

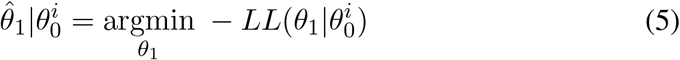

This is essentially optimizing *θ*_1_ for the minimum of *−LL* at different *θ*_0_ values, see Fig. 1D. The minimum of the PL is then subtracted, as the PL evaluates the likelihood ratio of different parameter sets with respect to the minimum *−LL*. The resulting PL *PL*(*θ*_0_) is a curve with minimum value at 0:

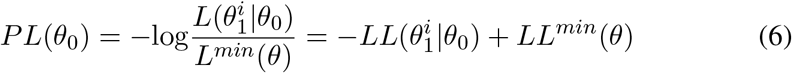

By Wilks’ theorem, the distribution of likelihood ratio asymptotically approaches *χ*^2^ distribution with increasing total number of cells [1, 2]. Therefore, the *χ*^2^ value (two tail: 1.92, half of 3.84 at one tail) at 1 degree of freedom at some significance level (*α*=0.05) was used to find the upper bound (*θ*_*ub*_) and lower bound (*θ*_*lb*_) for the CI on parameter *θ*_0_ from *PL*(*θ*_0_), as shown in Fig. 1E.

#### 3. Profile Likelihood for *a priori* identifiability analysis

We first study the identifiability of the telegraph model in the idealized, best case scenario: when the model itself is used to generate synthetic data. (We also call this ‘self-inference’). This is an *a priori* approach, because it depends only on the properties of the model itself and not on any particular dataset.

One can proceed directly from Eqn. 2, where *n*_*x*_ (the number of cells having *x* mRNA copies) is obtained from synthetic data, i.e., sampled from the CME-model-generated distribution (or, as is often done, simulated by SSA[3]). However, the likelihood then suffers from sampling error. This can be circumvented as follows: Let 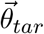 denote a target set of parameters, i.e., the set of parameters used to compute the ‘target’ distribution, which generates the synthetic data, and which we wish to infer. Then ⟨*n*_*x*_⟩, the expectation of *n*_*x*_, is 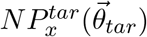, with *N* being the number of cells in this hypothetical experiment, and 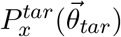 being the target distribution computed from the CME, as above.

In this way, Eqn. 2 can be further factored out the total cell number, *N*, giving

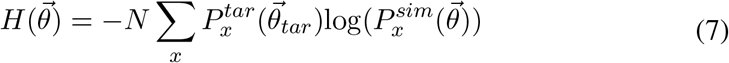

where 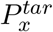 is the model-calculated probability of observing *x* mRNA in the target distribution, and 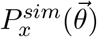 is the simulated probability of observing *x* mRNAs, given any parameter set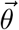. Eqn. 7 defines the cross-entropy between the target distribution 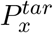 and the simulated distribution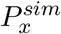, scaled by the number of cells *N*. It is minimized at 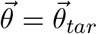, where it equals the scaled entropy of the target. The curvature near this minimum determines parameter identifiability: it corresponds to the Fisher Information Matrix (FIM), which bounds the variance of unbiased estimators. For consistency with the *a posteriori* analysis and standard likelihood terminology, we refer to the surface 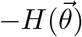 as the “log-likelihood surface,” while noting it is technically a scaled negative cross-entropy.

To study identifiability *a priori*, the question is what shape the hypersurface (Eqn. 7) has. In the ideal case, it is narrowly peaked, yielding relatively narrow CIs. However, if varying values of 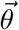 produce similar *P*, then the surface may have a broad peak with no clear global minimum–thus practically unidentifiable. The utility of Eqn. 7 is that one can study the effect of the cell number, *N*, in the hypothetical experiment on the *LL*, without any error introduced by sampling. CIs for each parameter can then be obtained from the *LL* surface using PL, as above.

We combine this *a priori* estimation of PLs (and hence CIs) with a computationally efficient method of globally scanning the parameter space, as a means to holistically study the identifiability of the parameters of the telegraph model.

Just as the *LL* hypersurface can be computed directly from *P* ^*tar*^ for an assumed cell number *N* without sampling (Eqn. 7), the PL can in turn be obtained in each parameter dimension without sampling. The blue curves in Fig. 2C,F are the PL obtained from ground-truth distributions, given an experiment with *N* cells. We refer to this as the ‘ground truth PL’. Each sample replicate (in red) can be considered to represent one experiment with finite (*N*) cells, whereas the ground truth PL is equivalent to the average of the sampled distributions’ PL, assuming an infinite number of experimental replicates with *N* cells, as illustrated here:

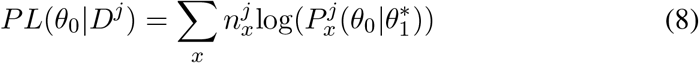

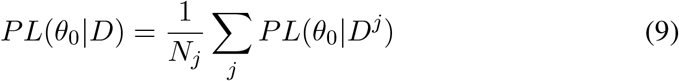

where *j* is the replicate/experiment index, *N*_*j*_ is the total number of experiments, and 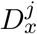 is the data sampled from the ground truth distribution in experiment *j*.

#### 4. Computational pipeline

##### Generation of the reference library from the CME

We use a simulated library of distributions as a reference to compute PLs for assessment of CIs/identifiability. To this end, To accelerate the computation, parameter sets were categorized by the maximum mRNA value based on *k*_*syn*_ so that the corresponding transition matrices generation and eigenvalue decomposition were performed batch-wise in numerical arrays, greatly reducing time for library generation. (For further details on how batch-wise/tensor format operations accelerate the library generation process, refer to SI 3.1). For the library used in this work, roughly 20 minutes were used in batch-wise/tensor operation, whereas it took roughly 2 hours in parallel in loop with an 8 core 16 threads laptop.

##### Strategy to combine coarse-grained library with fine-grained optimization

Obtaining accurate estimates of MLE and CIs involves non-linear optimization, which can suffer from local minima (termination point depends on initial guess), and the early termination problem (if the change in objective function is small per iteration because of a small gradient). It may also require many iterations to find the optimum if poor initial guesses and boundary conditions are given. This may not be a great concern if one is optimizing for one gene distribution, but it is not scalable when dealing with a large number of distributions, e.g. from transcriptome-wide data. To circumvent the poor initial value and boundary condition problems, our use of a coarse-grained, simulated library of distributions as a reference provides reasonable initial guesses and boundary conditions for optimization. Furthermore, the preliminary PL computed against the reference library provides a coarse-grained approximation. A larger/finer-resolution library provides more accurate coarse-grained PL curves and derived CI estimates, thus requiring less computation for latter optimization, at the cost of larger memory usage and time taken to generate the library.

The library is also used to rapidly assess whether further optimization is needed. Regions of the parameter space that are relatively insensitive to parameter changes have flat PLs, and thus wide CIs, and typically take more iterations for optimization. The preliminary PL, computed from these coarse-grained grid points, gives a rough idea of how identifiable each parameter is. Only distributions that are considered potentially identifiable would proceed to further optimization, under the rationale that precise quantification of very large CIs is generally not warranted, as long as these cases can be correctly labeled as ‘unidentifiable’. The criterion for further optimization is:

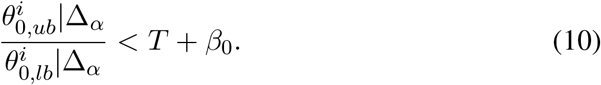

where 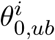 is the initial estimate of the upper bound on parameter 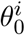, as determined by the coarse-grained PL (similar for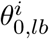). Δ_*α*_ is *χ*^2^ critical value, 1.92 at 1 degree of freedom, 5% significance level(*α*), two-tail. *T* is a threshold that depends on the order of magnitude of the feasible parameter range, and denotes an “acceptable” width of the CI on that parameter (see also next section). *β*_0_ is a buffer parameter that effectively makes the optimization criterion more generous. For example, it can serve to compensate for coarseness of the library grid, especially when the likelihood surface may have a steep gradient rendering initial CI estimates less accurate. As such, it is a user-defined parameter that can depend on factors such as library coarseness and sample size.

Following the same logic, if the optimization criterion is met, then the preliminary 2D likelihood plane also provides the boundary values and the initial guesses of the other parameter inputs for optimization. Optimization is performed within parameter bounds determined by the coarse-grained library as:

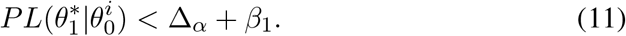

That is, optimization is performed within the region of parameter space where the initial library-computed PL is within Δ_*α*_ plus *β*_1_ of the minimum *−LL*. The value of *β*_1_ depends on cell size, as the log likelihood value is amplified by cell size, for larger cell number, a larger *β*_1_ should be used (-LL is far larger than Δ_*α*_). This potentially circumvents early termination at local minima if any, requires fewer iterations, and improves smoothness of the PL function in the vicinity of the MLE, promoting more accurate estimate of CIs. A non-linear optimization scheme, L-BFGS, was used to find the optimum. Though the *−LL* function does not have a specific form of gradient with respect to model parameters, using finite difference to approximate the gradient still allows quick convergence.

##### Profile Likelihood-based identifiability metric

In order to analyze the identifiability of the model itself, i.e., not specific to any assumed local parameter values (as in Fig. 2), but more globally over a broad region of feasible parameters, we first devised a PL-based metric for identifiability. A common measure is the Precision, defined as:

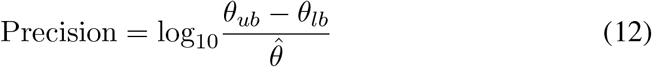

In general, the smaller the Precision, the smaller the CI and the more identifiable the parameter. We also defined another metric, termed the Alternative Precision Measure (APM):

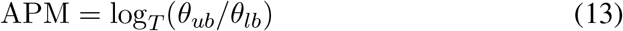

where *T* is a threshold value that is chosen depending on the assumed size of *θ*’s feasible parameter space. We used values of *T* for {*k*_*syn*_, *k*_*on*_, *k*_*off*_} = {3, 100, 100} (units of inverse degradation rate). The rationale for the different values is as follows: *k*_*syn*_ has a more linear effect on peak location, so the ratio between upper and lower CI bounds tends to be 𝒪 (1). In contrast, effects of variation in *k*_*off*_ and *k*_*on*_ tend to be more non-linear, and the parameters span several orders of magnitude. *T* sets the value of the ratio of the upper bound to lower bound at which *APM* = 1, i.e., an APM value of 1 for *k*_*off*_ indicates that *θ*_*ub*_ = 100*θ*_*lb*_.

Note that the APM is not scaled by 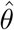 (unlike Precision). This ensures that a bad estimate of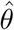, in cases of low practical identifiability, does not skew the result. In our testing, the APM provides a better measure of error as compared to Precision, since we are interested in quantifying large errors in order to fairly compare the practical identifiability of the model across different regions of parameter space. For further examples and discussion, see SI 2.1. APM will be the default identifiability metric of use, though we present results for both metrics.

##### Model for scRNAseq dropout

It is experimentally impossible to have 100% of the mRNA captured and amplified in scRNAseq, contributing to the “Dropout” problem of excessive zeros in scRNAseq[4]. To mimic such a process, our generated mRNA distribution library is modified with a binomial downsampling matrix with different capture rates [5], which are assumed known. Hence our modeling incorporates two types of technical error: limited sample size *N* and reduced capture rate (see SI 3.2).

**Table 1:**
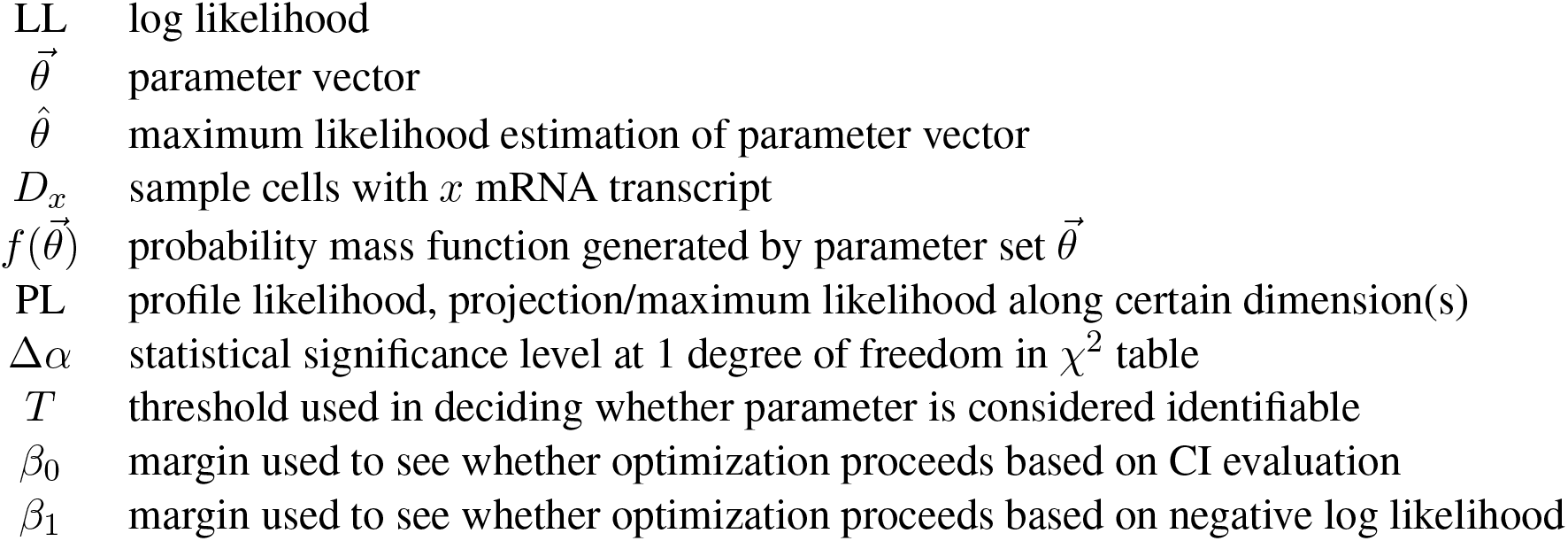
Variables and Metaparameters.

### 2. Extended Results

#### 1. Comparison of Identifiability Metric

We considered different measures of Precision, based on Confidence Intervals, in devising our identifiability metric. The Precision measure (Eqn. 12) takes the ratio between confidence interval (Upper bound-Lower bound) and MLE; since the binding(*k*_*on*_) and unbinding(*k*_*off*_) rates span over several order of magnitudes, we take the *log*_10_ of standard uncertainty. By this definition, accuracy of the MLE is higher if MLE is numerically larger. However, as the MLE strongly depends on noise, if we sample data from a distribution, the MLE can take place anywhere between the upper and lower bound, meaning as long as with the same confidence interval, MLEs with larger numerical value is equally good/bad as MLEs with smaller numerical value. This contradicts with the definition of Precision. Furthermore, as the three parameters span over different ranges of parameter space, it is inappropriate to compare identifiability of the different parameters in this way.

To circumvent this issue, we proposed a different metric that is independent of MLEs, but only dependent on confidence interval (APM, Eqn. 13). The threshold depends on the nature of the parameter, for example, for gene switching rates, they span over several order of magnitude, and their effects are more nonlinear, the threshold of choice is 2, meaning if the width of confidence interval spans 100, the switching rate is considered unidentifiable. For *k*_*syn*_, it spans over smaller order of magnitude, and its effect is more linear, the threshold of choice is 0.48, meaning the width of confidence interval spans over 3, the transcription rate is considered unidentifiable. Though one needs to decide what threshold to be used for each parameter based on the understanding of the model, nevertheless, it acts as a kind of normalization, and all parameters can be compared on the same ground. Though the two measures differ somewhat, there is broad qualitative similarity in their behavior over the parameter space. The resulting identifiability metrics of the parameter space is shown in Figure S1.

#### 2. Sensitivity analysis and comparison to distribution summary statistics

Here, the sensitivity vector of parameter *i* was computed as [6]:

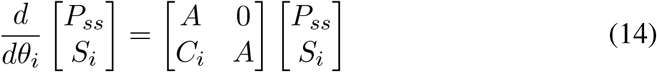

where *P*_*ss*_ is the steady solution to the CME, and *S*_*i*_ is the sensitivity vector of parameter *i. C*_*i*_ is the derivative of *A* with respect to *log*(*θ*_*i*_). *log*(*θ*_*i*_) is chosen over *θ*_*i*_ because the identifiability metric is based on the width of the CI which is in log10 scale. For more detail, please refer to SI 3.3.

The computed *S*_*i*_ is summed along axes other than those dimensions that are experimentally observed; here, the sum is over the off/on promoter states, since only mRNA copy numbers, but not underlying promoter states, are experimentally measured. Singular decomposition of the sensitivity matrix *S* reveals how sensitive the steady state distribution of the stochastic dynamical system is to the change of parameters. The smaller the minimum singular value, the less responsive the mRNA distribution to change of parameter values (with a zero value–singularity– indicating true structural non-identifiability).

**Figure S1:**
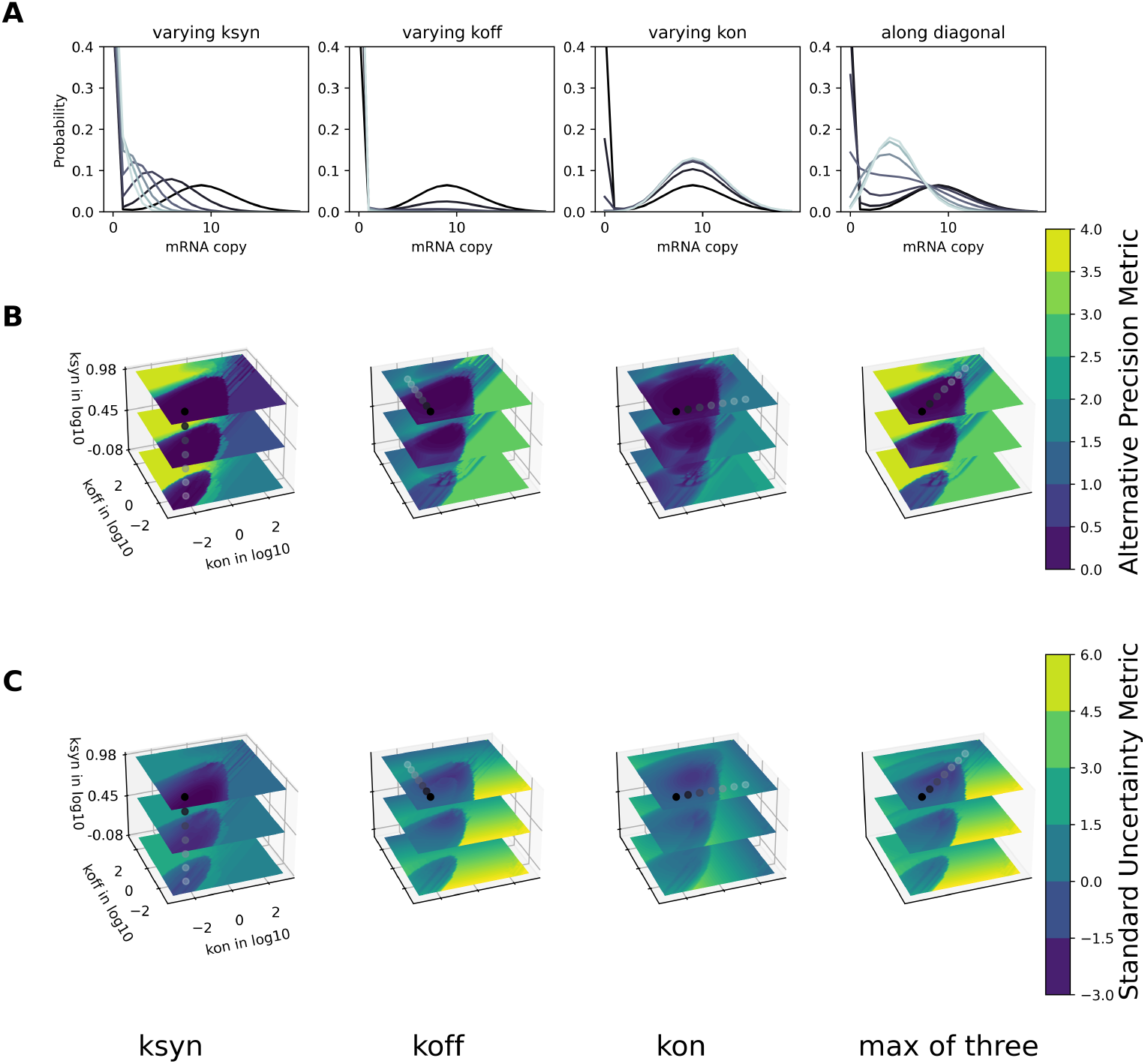
Comparison of the two identifiability metrics: A) mRNA distributions of parameter sets, top row with full capture rate; B,C) Identifiability of each parameter, and the max of three parameters as the overall identifiability of the parameter set for two metrics, Alternative Precision Metric and Standard Uncertainty Metric respectively.

We further explored how various RNA-distribution summary statistics varied as a function of the parameter values, reasoning that it could be useful if simple summary statistics (i.e., distribution shape measures) correlated with identifiability, thus providing an alternative means of *a priori* analysis. (Indeed, bimodality was seen to be a key feature, as described above). Two summary statistics that describe, at least in part, the distribution shape changes are the Fano factor *σ*^2^*/µ* (often used as a measure of dispersion in gene expression) and the ratio of the centralized third-order moment (denoted *M* 3) over the mean,

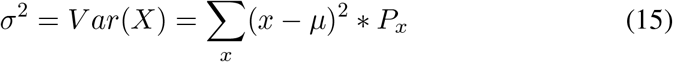

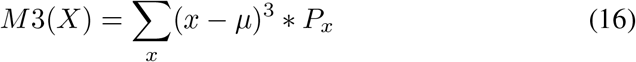

with *µ* the expectation value of the mRNA copy number. Division by *µ* ensures that Poisson-type distributions take the value of 1 for both metrics. 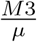 works like a higher order analog of the Fano factor, correlating with skewness of the distribution rather than dispersion. Positive values indicate skew towards low RNA copy numbers.

**Figure S2:**
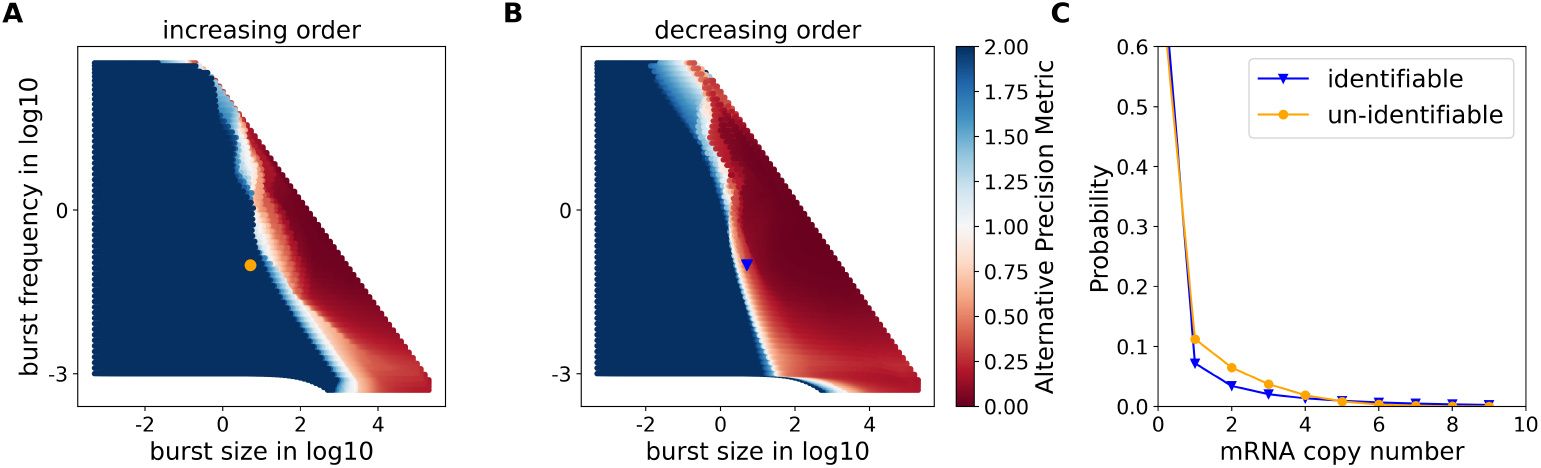
Identifiability of transcriptional burst-size and burst-frequency via the telegraph model. The two representative parameter sets have the same burst dynamics (burst size: 5.13, burst frequency: 0.0975).(A,B) Scatter plots of APM (maximum over all three model parameters) from self-inference for 1000 cells plotted versus burst-frequency and burst-size. Information from 3D {*k*_*on*_, *k*_*off*_, *k*_*syn*_} (data as in Fig. 3) is projected onto the 2D space {burst-size, burst-frequency}. (There is a nonlinear relationship between the latter and the former, see Methods). Different projection methods are shown: (A) Increasing order, means that for a given point in the 2D plane, which maps to multiple points (computed APM values) in the 3D space, the highest APM value is visualized. (B) Decreasing order, means that the lowest APM value is visualized. Highlighted points correspond to two representative parameter sets, (*k*_*syn*_ : 20, *k*_*off*_ : 3.9, *k*_*on*_ : 0.1, orange point in (A)), (*k*_*syn*_ : 2.67, *k*_*off*_ : 0.52, *k*_*on*_ : 0.12, blue point in (B)). These two sets have the same burst size and burst frequency, but APM values of 1.86 (un-identifiable) and 0.448 (identifiable), respectively. (C) mRNA distributions of the two parameter sets, showing similar but different shapes.

Both *M* 3*/µ* (Fig. S3B) and Fano (Fig. S3C) generally have larger magnitudes in the inverted cone region of the parameter space where APM shows better parameter identifiability. Both measures take values close to 1 when either *k*_*off*_ ≫ *k*_*on*_ (gene mostly off, high cell number with 0 mRNA) or when *k*_*on*_ ≫ *k*_*off*_ (gene mostly on, Poisson distribution). This indicates that, in general, distributions with complex shapes (higher dispersion and/or skewness) can be predicted to have more identifiable bursting kinetics. However, both measures have limitations. For example, the Fano factor takes values close to 1 in the region of slow *k*_*off*_ and medium *k*_*on*_, although there is moderately good identifiability in this region. This is because the distribution is bimodal, though with low probability in the 0-mRNA peak, which contributes little to the variance. *M* 3*/µ* performs somewhat better in this same region, but is challenged when distributions are close to symmetric (*k*_*on*_ *≈ k*_*off*_), albeit bimodal. When *k*_*off*_ ≫ *k*_*on*_ (distribution nearly unimodal at 0 mRNA, but with a small second peak affecting dispersion and skew while *µ* is low), both metrics can take high values despite rapidly decreasing identifiability in this region as one exits the inverted cone region. Additional summary statistics were also studied (Fig. S4).

**Figure S3:**
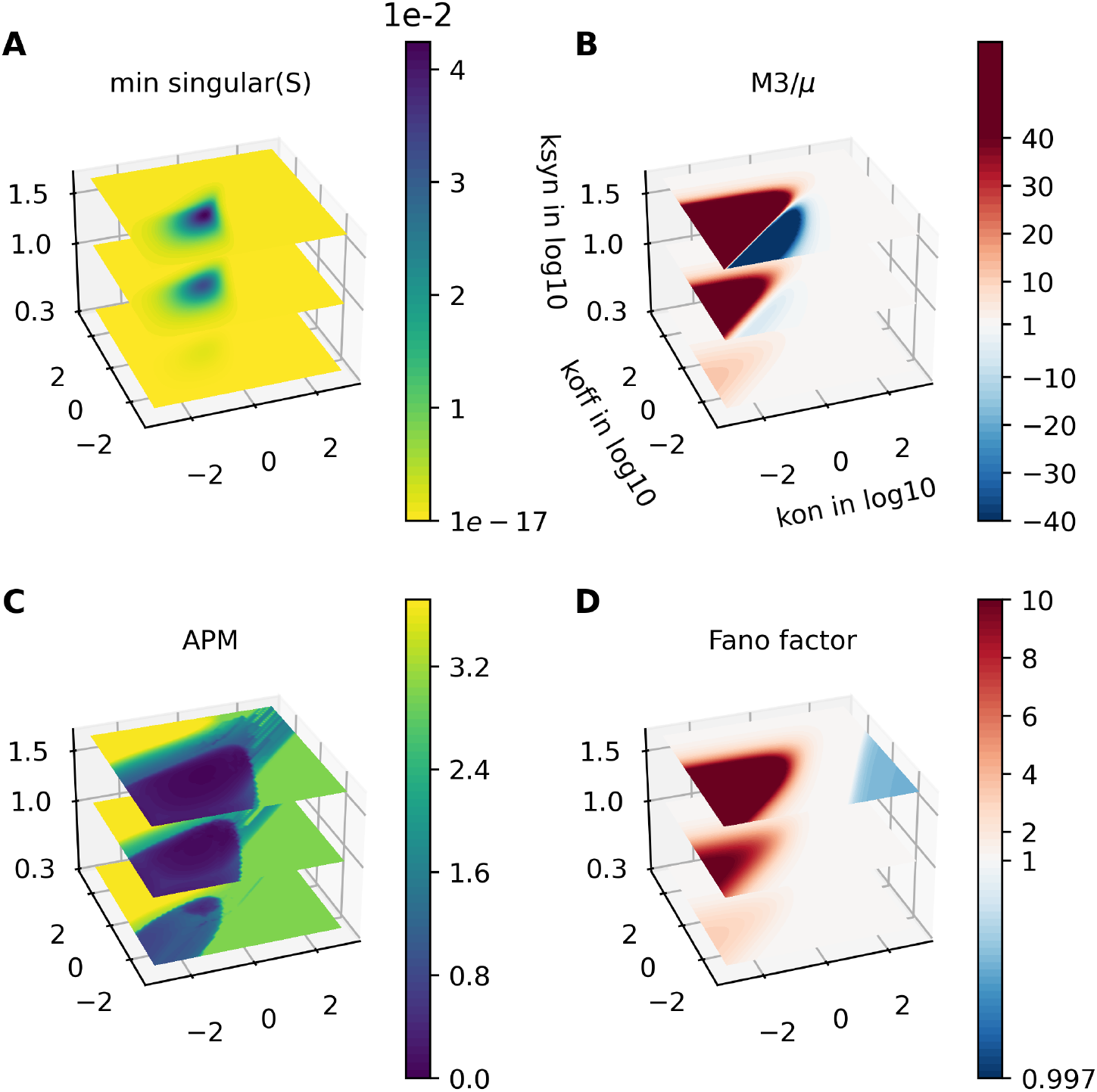
Comparison of different approaches to *a priori* identifiability analysis over the parameter space. (A) Minimum singular value of the sensitivity matrix *S*, derived directly from the CME model; (B) Summary statistic *M* 3*/µ*, the centered 3rd moment divided by the mean; (C) Profile-Likelihood-based identifiability for comparison (maximum of the APM for 1000 cells and 100% capture, same as Fig. 3B, right); (D) Summary statistic Fano factor. Note that for B and D, a different color scale centered on 1 is used, to better highlight regions of non-Poisson distribution shape.

### 3. Neural Net-based prediction of parameter identifiability

Different distribution statistics measures were used as neural network input features. The list of statistics measures includes: Fano factor, mean (*µ*), variance (centralized second moment, M2), centralized third moment (M3), 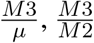. Some of the statistic measures are visually more similar to the identifiability of parameter space computed from the computation pipeline in some region, but not entirely matching. Fig. S4 shows the different measures.

No single shape metric studied correlated perfectly with identifiability as quantified through the PL-based pipeline We reasoned that an identifiability predictor could be trained on our self-inference results, together with information on summary statistic features. Since identifiability is related to distribution shape, we asked whether it would be possible to rapidly assess identifiability for a given experiment-derived mRNA distribution (together with information on cell number and capture rate) without applying the full PL-based inference pipeline. In practice, only distributions predicted to be identifiable would then proceed downstream in the pipeline. This would reduce the amount of optimization required. However, with just summary statistics like moments and Fano factor, it is impossible to fully separate parameter sets with low and high APM at different capture rates and sample size with a simple linear regression model. To leverage the powerful generalization capability of deep learning, a neural network was trained to predict whether a distribution is identifiable or not. Summary statistics (those of Fig. S3 and Fig. S4) and sample size were used as input, and the APMs were used as output. The simulated data (60^3^ distributions, at five sample sizes:10^2^, 10^3^, 10^4^, 10^5^) were used for training and testing at 7:3 ratio, respectively (training curve shown in Fig. S5A, data shown in Fig. S5 is from the deep learning model with the simplest multi-layered fully connected neural-net). When parameter sets were classified (identifiable, non-identifiable, very non-identifiable), 96.7% of identifiable parameter sets (in ground truth) were also predicted to be identifiable (Fig. S5B). Deeper analysis into the remaining sets that were wrongly predicted to be un-identifiable revealed that, in these cases, summary statistics used to train the neural net were not sensitive to small details in distribution shape that nevertheless promote identifiability and are somehow utilized in the PL-based pipeline, e.g., bimodality characterized by extremely low probability in the second (high copy number) peak, and bimodality that is degraded by down-sampling (i.e., to model low capture rate), but still gives the distribution a slightly modified shape. (See also discussion related to Fig. S3). Predictive capability of the neural net for the identifiability of each individual kinetic parameter is shown in Fig. S5C-E.

**Figure S4:**
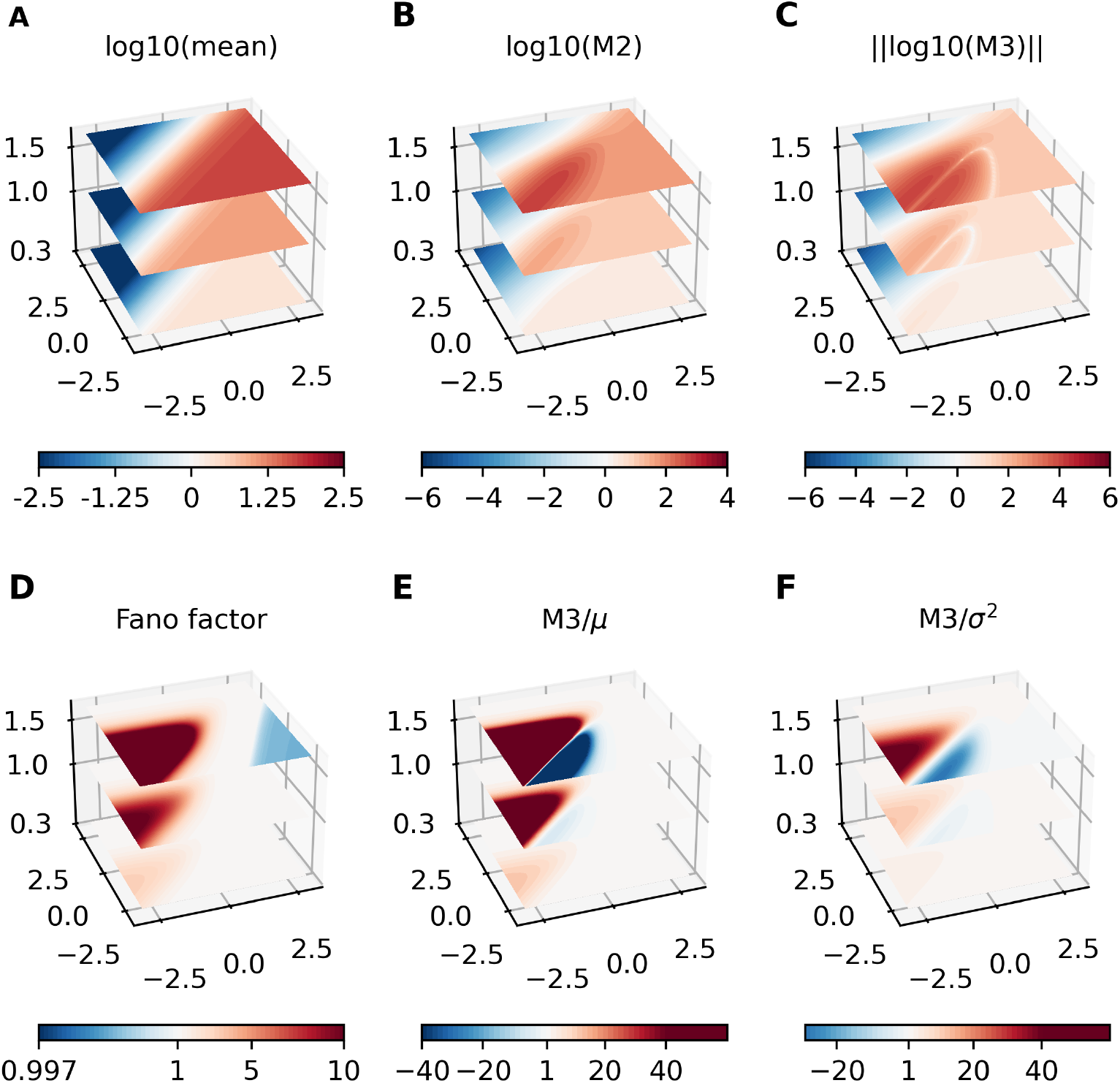
Comparison of different statistical metrics of the parameter space: A) Mean (*µ*) in log10, B) Variance(M2) in log10, C) absolute value of centralized third order moment (M3) in log10, D) the Fano factor, E) 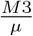, F) 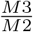.

The neural network predicted 1945 genes in the SS3 data to be identifiable (that is, roughly twice as many genes than were truly identifiable, according to the PL pipeline). However, there was good overlap (877/970) between the set of identifiable genes (via PL) and those via the neural network. Similarly, for HUES64WT data sets, the neural network predicted 1384 genes to be identifiable, and 967 genes were considered identifiable according to both methods. Deeper analysis of the genes that were falsely predicted to have identifiable kinetics, according to the neural network, revealed that it is due in part to distribution features found in the real-world datasets that are under-represented in the simulated distribution library (namely, low mean and variance, whereby most probability lies at 0 mRNA with low probability of low copy number expression). The majority of these distributions have predicted APM between 1 and 2.5. These distributions mostly locate near the boundary of identifiable and un-identifiable region in Fig. S3.

**Figure S5:**
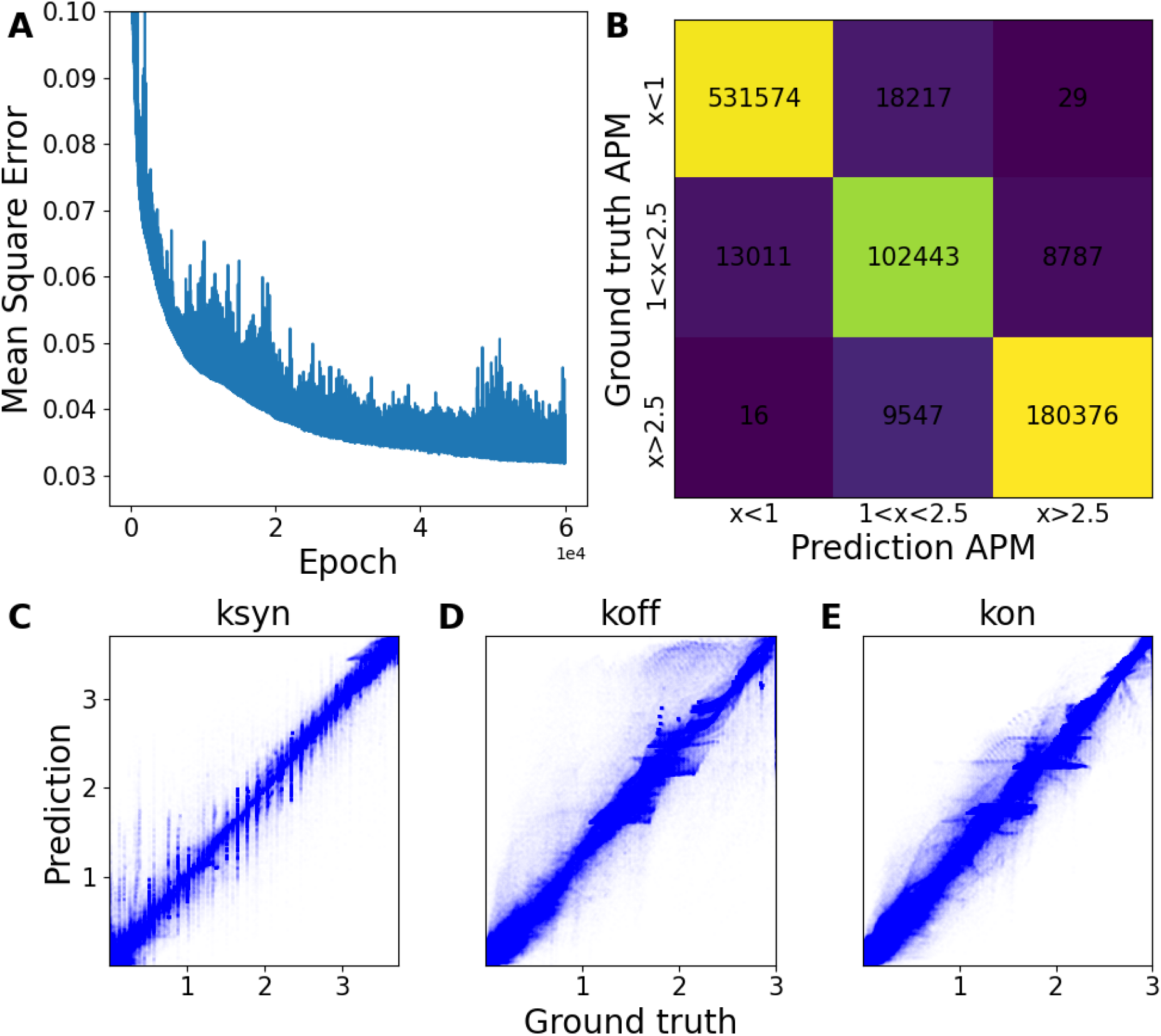
Neural network prediction of parameter identifiability (as quantified by the APM) based on sets of statistical features or RNA distributions at 30% capture rate. A) Training curve of the neural network with testing mean square error; B) Confusion matrix for prediction of APM against ground truth. The number represents number of parameter sets. Larger values on the diagonal indicate better prediction. Parameter sets were classified into identifiable (*x <* 1, where x is the APM value), practically un-identifiable (1 *< x <* 2.5) and very practically un-identifiable (*x >* 2.5); C-E) Scatter plots of predicted APM value for each individual parameter over the full simulated library, versus the ground truth APM value. A good prediction should have the majority of dots near the diagonal.

### 3. Supplementary Methods

#### 1. Transition Matrix Dimension Reduction

Gene switches between active (*G*^*\**^) and inactive(*G*) stochastically with reaction rates *k*_*off*_ and *k*_*on*_ respectively.

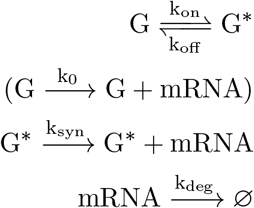

The basal expression rate *k*_0_ is optional depending on the assumption. (1) If we are only interested in telegraph model: we can set basal expression to be null (*k*_0_=0); (2) In the case of non-zero basal rate, with some basal expression rate (*k*_0_ ≠ 0). *k*_*syn*_ = *k*_0_ *\* k*_*m*_, where *k*_*m*_ represents the binding effect of the transcription factor.

Let *P*_*k*_ be a vector of probability at mRNA state *x*_0_,…,*x*_*n*_ at gene state k. The above system can be written in block matrix form:

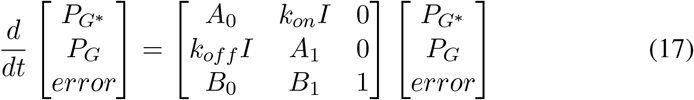

where I is the identity matrix. The error term absorbs all the degradation and production probability from the two boundary at *x*_0_ and *x*_*n*_ respectively. The matrix *A*_*k*_ is a tridiagonal matrix denoting the connections within states in *P*_*k*_:

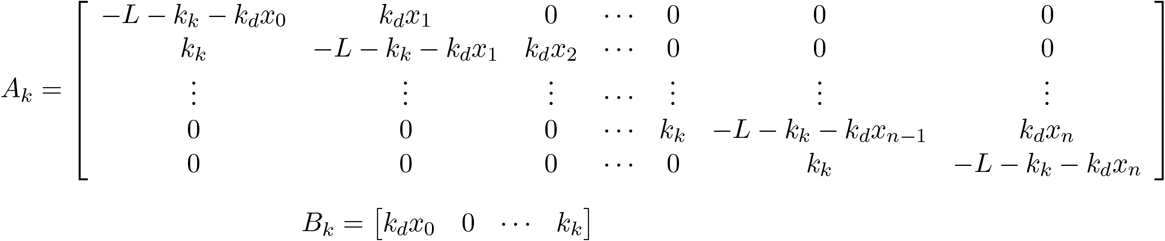

where *k*_*k*_ is the transcription rate of at gene state k, and *L* is *k*_*off*_ if *A*_*k*_ is *A*_0_, or *k*_*on*_ if *A*_*k*_ is *A*_1_.

At steady state, left hand side of Eqn.17 is 0. Instead of solving the SVD of the whole transition matrix, for Eqn.17, it is rather convenient to solve it as linear system in block matrix form.

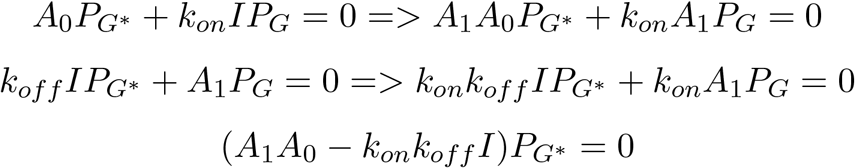

applying SVD on 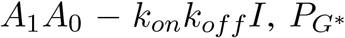 is obtained. However, it needs to be normalized and scaled. Unlike directly applying SVD on Eqn.17, the steady state distribution only needs to be normalized to 1. 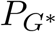 needs to be further scaled by the probability of gene state *G*^*\**^, which can be obtained from below:

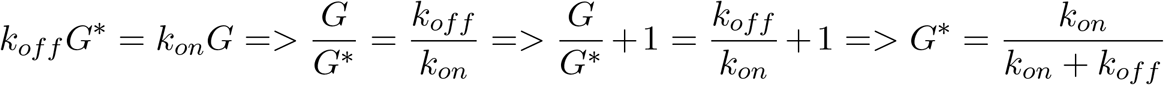

Once 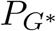 is obtained, *P*_*G*_ can be easily calculated. By doing in block fashion, solution is obtained by SVD on a matrix of size n, instead of SVD on a matrix of size 2n. This would largely increase computation efficiency, as the time complexity of SVD is *O(n*^3^). Fig. S6a illustrates how time increase in SVD changes against the matrix dimension. The increase in time grows exponentially as system dimension increases. The time taken for SVD operation are similar for operation in batch/tensor and in loop. However, significant difference in construction of the transition matrices between the two approaches. Fig. S7a demonstrate the accuracy of the solution compared to the one for solving full size transition matrix *A*. Since for a simple two-state switching system, there is no inverse calculation, there is practically no difference. Fig. S7b demonstrate the computation speedup by solving in blocks.

In theory, regardless the number of gene states, the system can be reduced to SVD of matrix size n, with some linear algebra. For example, considering two promoter regions where two transcription factors can bind to, leading to four different gene states. Again, one can put different assumptions in the model. For example, (1) if both the promoter regions are bound, if there exist inhibitor, inhibitor wins, otherwise, the *k*_*ab*_ takes the larger value of *k*_*a*_ and *k*_*b*_; (2) *k*_*ab*_ is the sum of *k*_*a*_ or *k*_*b*_, etc.

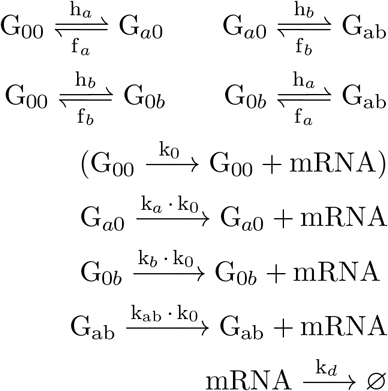

**Figure S6:**
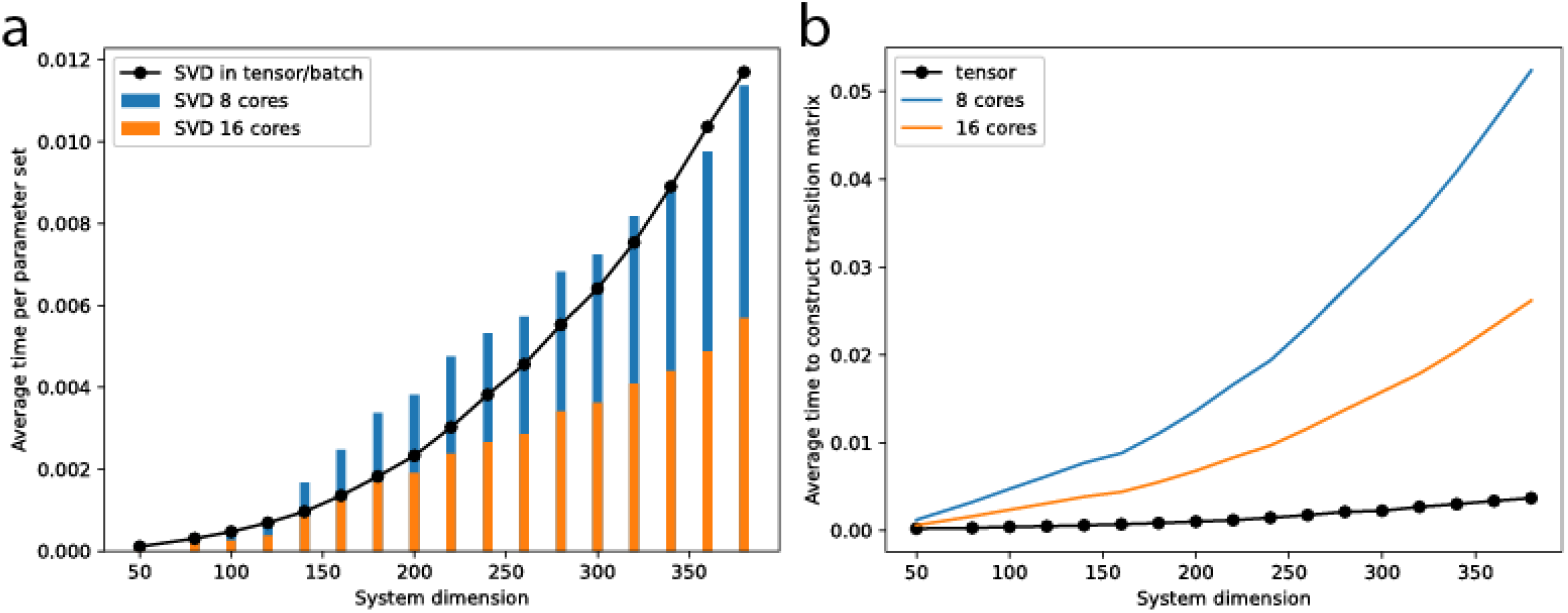
(a) Computation efficiency of SVD at different system size and (b) Efficiency in constructing transition matrix in tensor/batch by a 8-cores,16 threads configuration; The blue and orange bars are operation in loop wise by a 8-cores,8 threads and 16-cores,16 threads setting running in parallel; The black dotted line is operation in batch-wise/tensor format in 16 threads setting.

Again, the system can be represented in block matrix fashion:

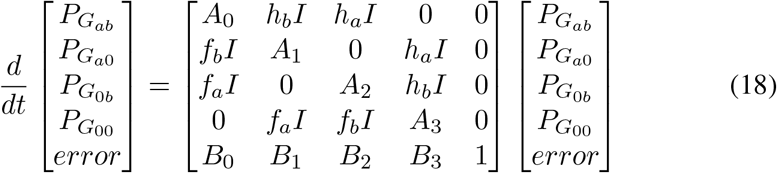

**Figure S7:**
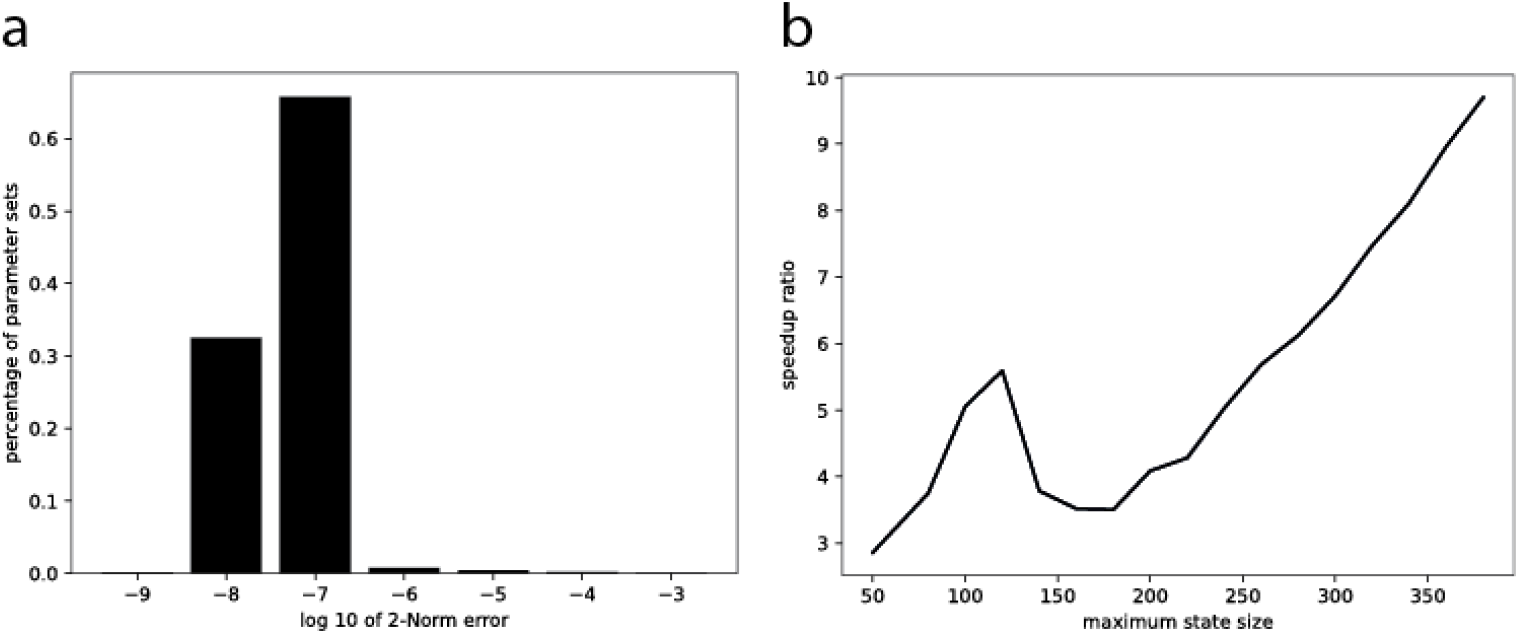
Error and speedup comparison for two-state model;(a) Histogram of 2-norm error over the parameter space between solution of full-transition versus the one of block transition; (b) Speedup times of solving SVD on the same system with block approach relative to full-transition approach

Similarly, one can solve (18) block-wise with some linear algebra:

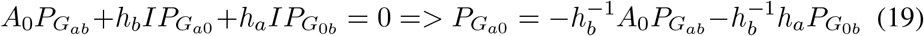

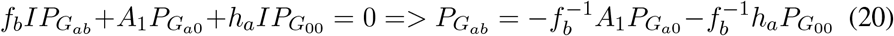

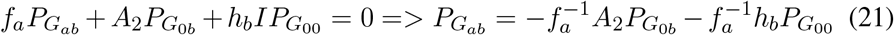

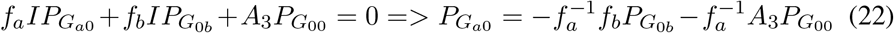

(19) = (22):

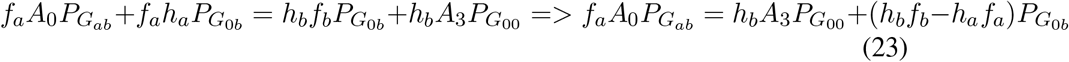

Plug (21) to (23):

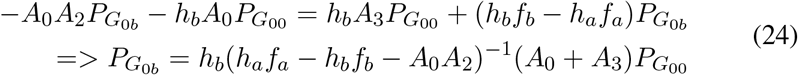

(20) = (21):

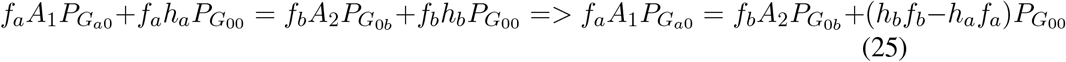

Plug (22) to (25):

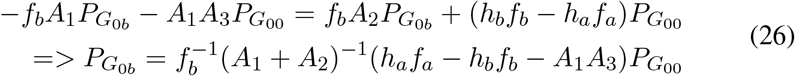

(24) = (26):

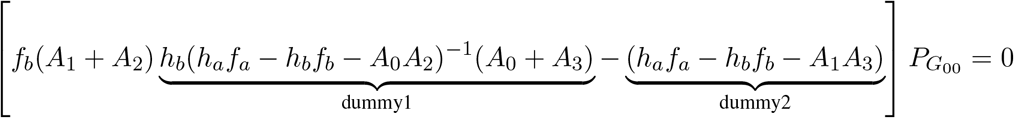

**Figure S8:**
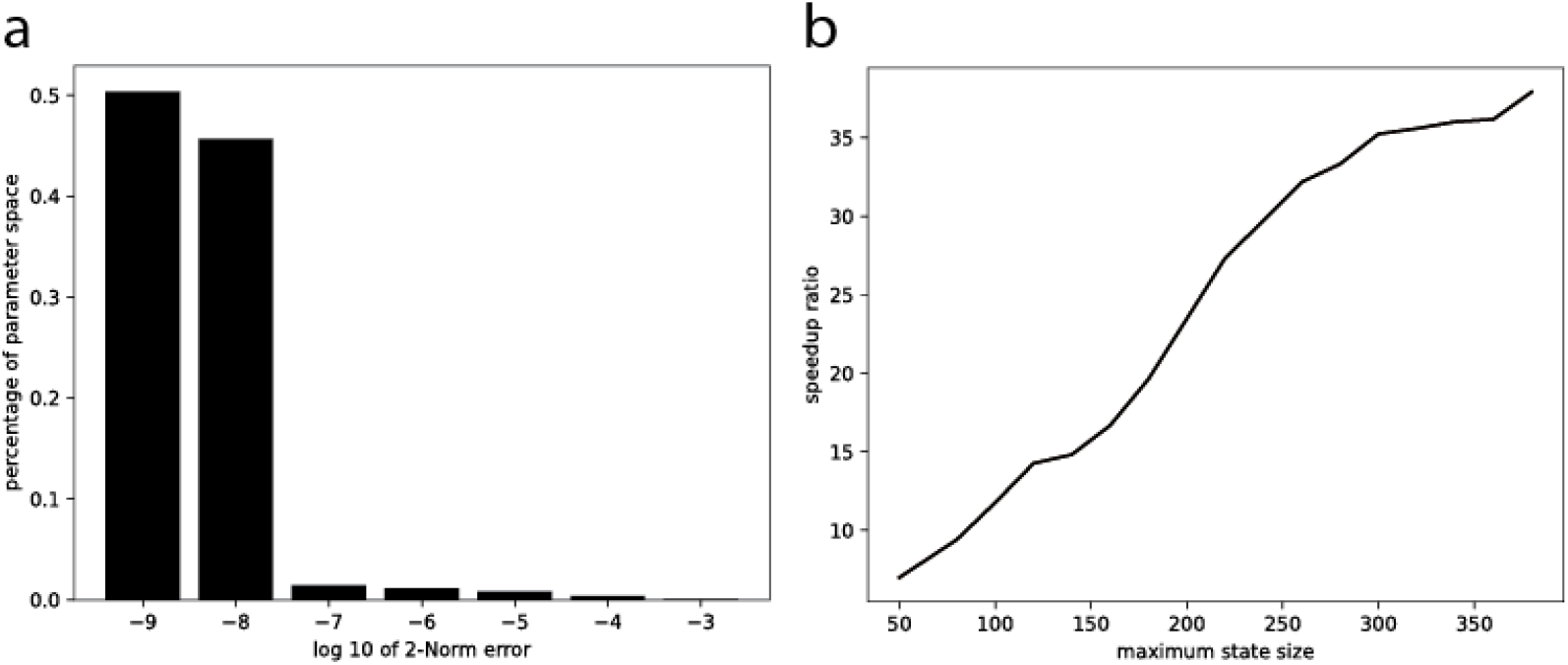
Error and speedup comparison for four-state model;(a) Histogram of 2-norm error over the parameter space between solution of full-transition versus the one of block transition; (b) Speedup times of solving SVD on the same system with block approach relative to full-transition approach

Again, 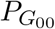 is obtained by performing SVD, and scaled to have sum with probability of state *G*_00_:

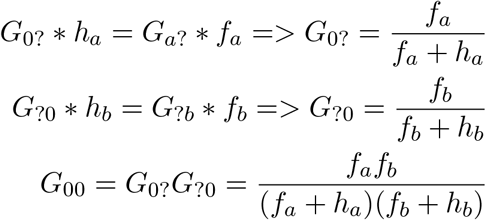

The rest of the probability distribution at other states can be calculated accordingly. Overall, for one-gene four-state model, one inverse matrix of size n is calculated and one SVD on matrix of size n is performed. As long as the transition matrix is written in the form of block based, and system states are categorized by gene states, the result of CME can be computed in a reduced form. It is much more efficient memory-wise and significantly faster compared to performing SVD on Eqn.18, matrix with size n*number of gene state. Fig.S8a shows the accuracy of the solving four-state system in block-wise compared to full transition matrix *A*. Fig.S8b demonstrates the computation acceleration.

To make use of efficiency of vectorized operation in modern computing, we put the transition matrix A of different parameter sets into a tensor format, and make all the algebra calculation batch wise. This would lead to significant speedup compared to loop-wise, so that the parameter space can be explored comprehensively and efficiently. The reduction in time is twofold: (1) Well-built package for tensor operation utilizes computation resources more efficiently(hyper-threading, multi-cores); (2) Vector operation is faster then looping. Fig.S6a shows the computation acceleration by performing SVD in tensor(batch) format. For system dimension up to about 340, performing in tensor/batch is no slower than in loop if not faster.

Though one can construct transition matrix by evaluate the minimum number of states from every set of kinetic parameters, this would also lead to significant overhead time. In contrast, by first categorizing the parameter sets into different group of maximum dimension systems, then constructing transition matrix batch-wise can lead to significant lower overhead computation, as shown in Fig.S6b.

### 2. Mimicking Low Capture Rate in Sequencing

For inference of experimental data, since it is impossible to have 100% of the mRNA captured and amplified in scRNAseq, the actual mRNA distribution is downsampled with a certain capture rate, leading to an observed distribution with excessive zeros. To mimic such a process, the generated mRNA distribution library is modified with a binomial downsampling matrix with different capture rates[5]. Hence our modeling incorporates two types of technical error: limited sample size *N* and reduced capture rate. One can describe such binomial downsampling process as *Bi*(*i, x*), the probability of observing *i* mRNA when there are *x* mRNAs in cell in a binomial distribution, and the observed/downsampled mRNA distribution can be found as:

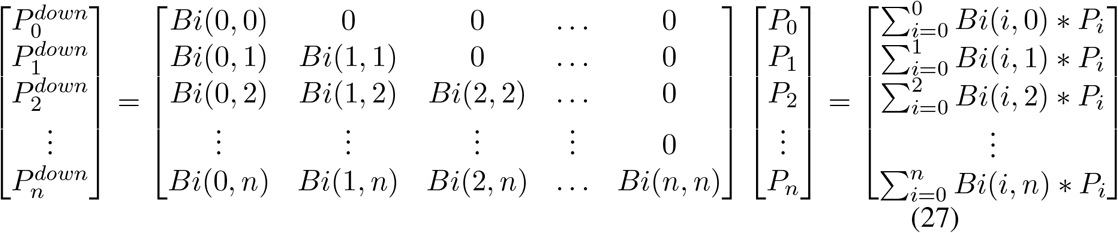

Here, we assumed that mRNAs from different genes share the same probability to be captured/amplified during sequencing process.

### 3. Sensitivity Computation

In the context of CME, we can compute sensitivities with a little modification (constraint of sum of 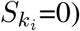:

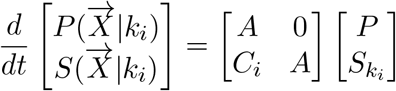

where 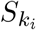 is the sensitivity of model output with respect to parameter *k*_*i*_, and *C*_*i*_ is the partial derivative of *A* with respect to parameter *k*_*i*_. Sensitivity matrix only represents the local effect, however, the effect of each parameter on the resulting mRNA distribution in non-linear. Further, identifiability of parameter set is quantified by the width of confidence interval in log10 scale, therefore, instead of directly calculating sensitivity with respect to parameter value, a better approach is to calculate with respect to the log of parameter:

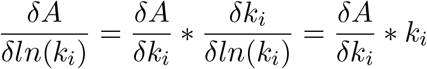

With a known 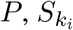 is obtained relatively easily with condition that sum of 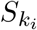 is 0 (total change/flow of probability is 0). One can construct the sensitivity matrix from the sensitivity vector respective to each parameter, and by performing SVD, the smallest singular value and its corresponding unitary vector represent the magnitude and direction where each parameter reflects to the minimal change to the corresponding parameter/least sensitive. In stead of using the dimensionless sensitivity where the sensitivity vector is normalized by 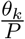, we kept the absolute value here. A comparison of the minimum singular value of sensitivity with respect to parameter value and the log of parameter value is shown in Figure S9.

---

**Figure S9:**
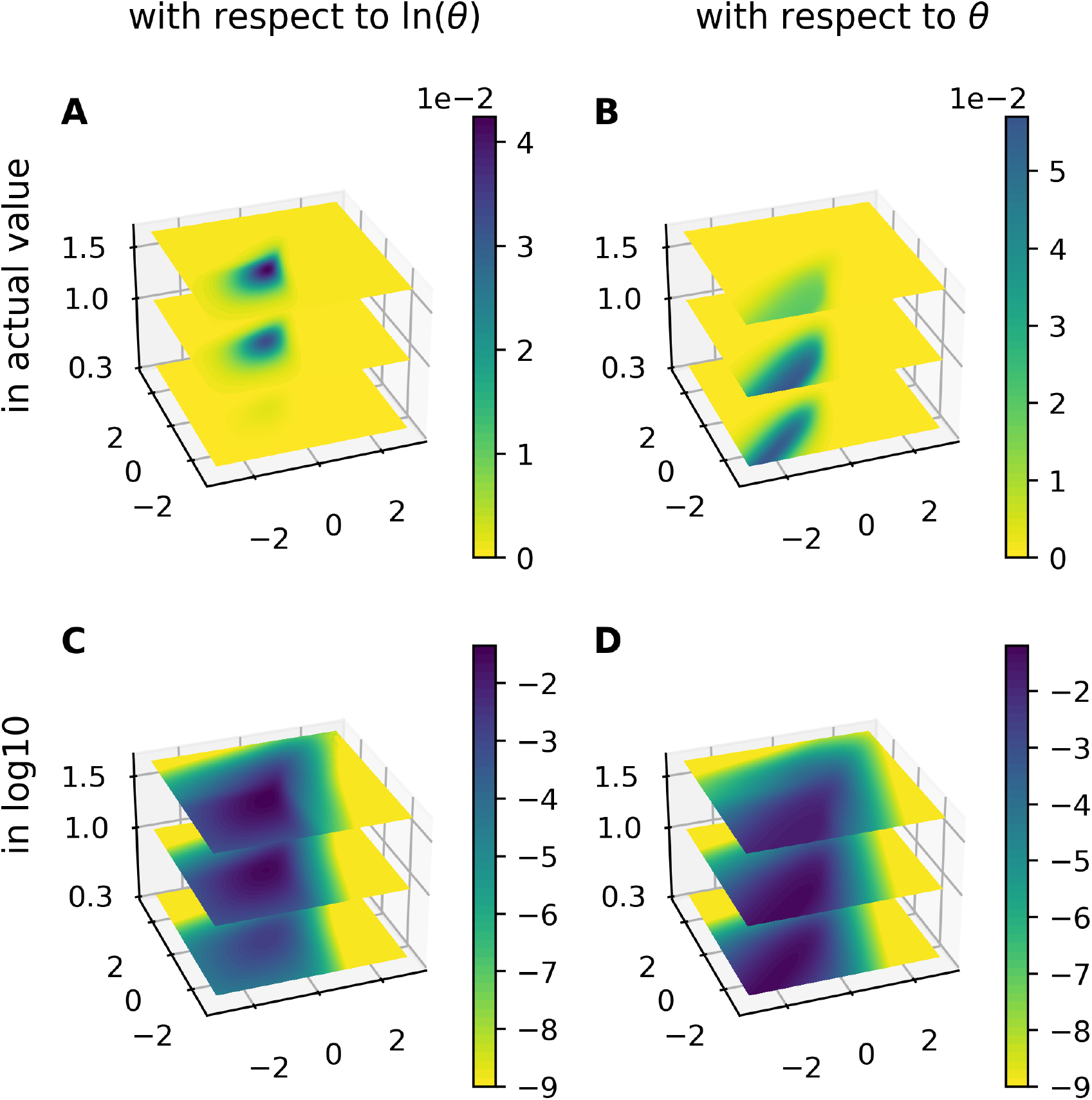
Comparison of minimum singular value of sensitivity with respect to log of parameter (first row) and with respect to parameter (second row), the log10 of singular value are shown in the second column, to better shown the order change of singular value

## Notes

### Competing Interest Statement

The authors have declared no competing interest.

